# Spatial proteomics with 100+ markers with Highly Multiplexed MALDI-IHC

**DOI:** 10.1101/2025.04.18.649415

**Authors:** Mengze Zhang, John Abbey, Pierre Bost, Genki Usui, Simon Häfliger, Simone Muenst, Mark J. Lim, Gargey B. Yagnik, Natalie De Souza, Kenneth J. Rothschild, Bernd Bodenmiller

## Abstract

Highly multiplexed, antibody-based protein imaging of tissue has yielded many insights but does not yet allow the routine analysis of more than 50 markers simultaneously. By leveraging the ability of Matrix-Assisted Laser Desorption/Ionization (MALDI) imaging to resolve hundreds of m/z peaks without spectral overlap, MALDI-IHC with photocleavable mass-tag (PCMT)-labelled antibodies offers the possibility to dramatically increase the number of markers that can be imaged. Here, we demonstrate 100+ plex MALDI-IHC imaging and describe our development of gutenTAG, an open-source analysis pipeline for the resulting highly multiplex data. We validated a panel of 133 tumor-ecosystem-focused antibodies, show that it captures expected histopathology in diverse tissues, and demonstrate reproducibility of the method. We further applied a 120-plex subset of this panel to samples from an endometrial cancer cohort, again capturing expected histopathology and identifying novel spatial structures between TCGA molecular subtypes. Our work demonstrates the feasibility of simultaneously detecting hundreds of protein markers in a reproducible and scalable manner.

## Introduction

Biological systems are complex and require sophisticated analytical tools for comprehensive visualization and understanding. Spatial omics technologies have emerged as powerful methods for depicting biological systems via profiling of transcriptomes, proteomes, metabolomes, and of other molecular classes such as glycans and lipids. Multiplex antibody-based protein imaging probes the expression of multiple markers in tissues at cellular or even subcellular resolution and is now applied in both the research and increasingly the translational setting^1–5^.

However, the routine multiplexing capacity of protein imaging is currently limited to around 50 markers per analysis^6,7^. Metal-isotope based methods like imaging mass cytometry (IMC) and multiplexed ion beam imaging (MIBI) are limited by the available isotopes. Cyclic staining^8,9^ or DNA barcoding^10–12^ using immunofluorescence (IF)-based methods can further increase the plexity and has been reported to achieve over 100-plex imaging^13,14^, but cyclic methods are limited by lengthy protocols, increased background signals over time, and sample degradation with potential tissue morphology changes over imaging cycles. Alternatively, sequencing-based methods such as CITE-seq^15^ also enable high-plex proteomics imaging, but require subsequent sequencing of DNA barcodes on antibodies, and are thus both time- and resource-intensive.

One solution to drastically increase the multiplexity of single-round protein imaging is to employ photocleavable peptide mass tag (PCMT)-labelled antibodies^16^, termed MALDI-IHC^17^. Rather than labelling antibodies with metal isotopes as mass-tags in (IMC)^6^ and (MIBI)^7^, antibodies for MALDI-IHC are labelled with PCMTs followed by imaging with matrix-assisted laser desorption/ionization time-of-flight mass spectrometry imaging (MALDI-TOF-MSI). Given the ease of design and synthesis of a peptide mass tags and the analytical performance of MALDI-TOF instruments (i.e., high mass resolving power and mass detection range, spatial resolution ≥ 5µm)^18–22^, MALDI-IHC has the potential to reach >100-plex protein imaging at near-single-cell resolution in a single staining and measurement step. The ability to achieve high plexity is also enhanced by the design of novel PCMTs which incorporate a photocleavable linker which exhibits a highly efficient and fast photoreaction and after photocleavage the bulk of the photocleavable linker remains attached to the antibody. In addition, the photoreleased mass reporter has a positive charged amino group which enhances detection of these positively charged ions^39^.

Proof-of-principle work using MALDI-IHC has so far shown the simultaneous imaging of only 12 markers in human formalin-fixed paraffin-embedded (FFPE) tonsil and breast cancer samples^17^, and using a 27-plex panel 15 markers were imaged in combination with lipids MSI on the same sample from a fresh frozen breast cancer sample^39^, thus the full potential of this technology remains unexplored. Further, existing MALDI-imaging data processing tools are designed for untargeted datasets, often relying on high-mass resolution instruments for precise peak annotation. In contrast, MALDI-IHC is a targeted approach that does not require complex peak annotation algorithms but faces specific challenges like peak widening and the absence of quantitative quality control metrics. The lack of an open-source pipeline tailored for high-plex MALDI-IHC across different instrument types is a major barrier to efficient data analysis.

In this work, we developed and validated a 133-plex PCMT-labelled antibody panel for MALDI-IHC imaging across multiple FFPE tissue types, assessing reproducibility in whole-slide imaging of serial sections and using parallel histological annotation to validate the identified tissue regions.

For data analysis, we developed *gutenTAG*, an end-to-end and open-source preprocessing, quality control and data analysis pipeline in R. Using a 120-plex panel focused on peripheral solid tumors, we analyzed samples from 18 endometrial cancer (EC) patients, identifying shared and molecular subtype-specific spatial features but also substantial intra- and inter-patient heterogeneity. Overall, we demonstrate the feasibility and reproducibility of 100+ plex protein imaging, capturing key histopathological features across diverse tissue and disease types.

## Results

### A panel of 133 validated antibodies for MALDI-IHC

To develop ultra-highly multiplexed protein imaging with MALDI-IHC, we assembled an antibody panel comprising 133 antibodies designed to characterize the tumor microenvironment (TME) with a focus on solid tumors. We selected antibody clones that had been validated for IMC from previous studies^34–38^, targeting phenotypic markers for tumors (25), tumor microenvironment components (9), metabolism markers (7), immune cell markers (43), pathway markers (16), cell state markers (10), and disease-specific markers (10), and further incorporated 10 neurology-related markers (**Table 1**). Each antibody was labeled with a unique PCMT of a defined mass over charge (m/z) using an amine-NHS click chemistry approach^17^ (**Fig. 1a**). The PCMTs used were designed to have similar ionization efficiencies^17,39^ and were therefore randomly assigned to each antibody. After validation and antibody titration, these labeled antibodies can be employed individually or combined to create multiplexed staining panels (**Fig. 1b**) since there is no overlap between the detection channels.

**Table 1.**
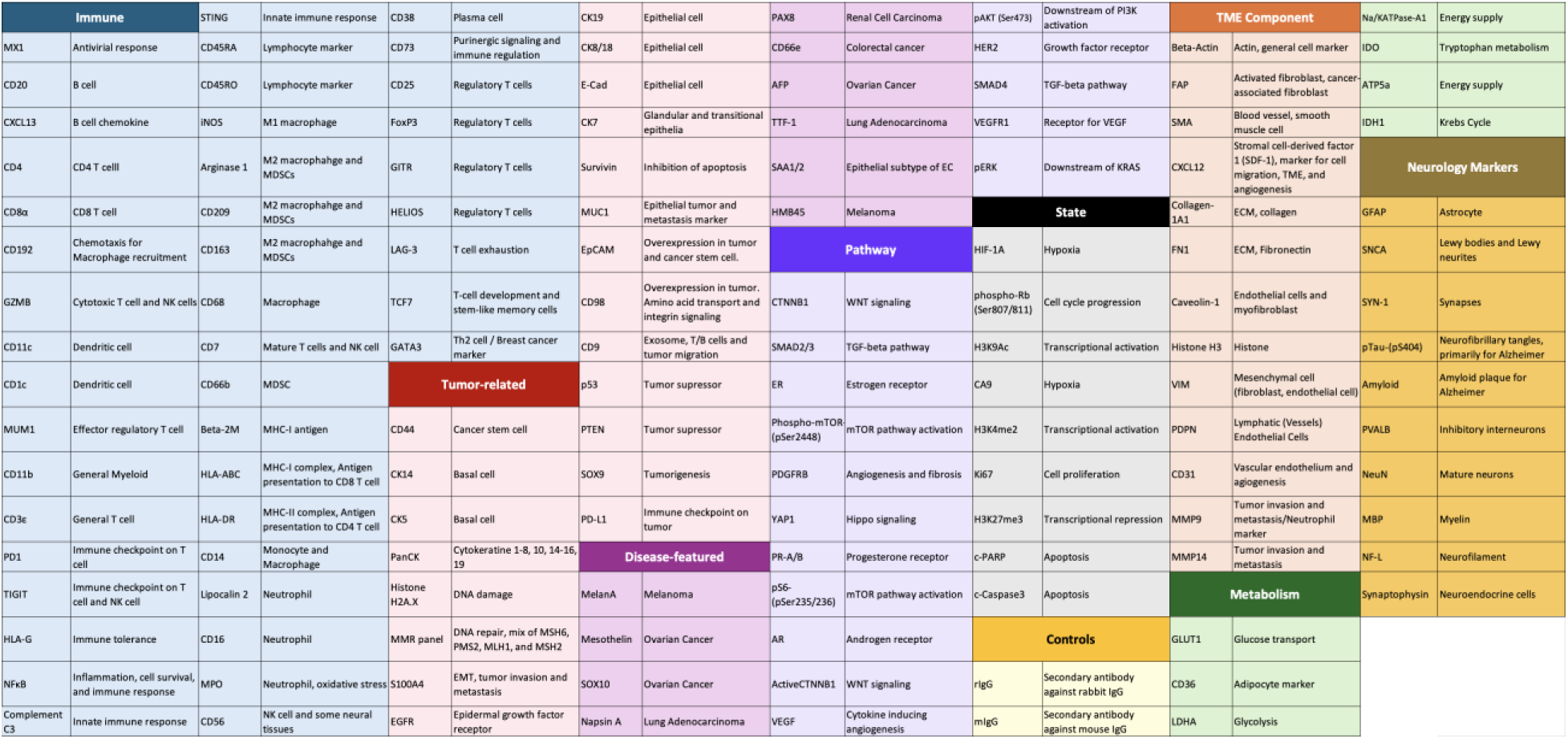
Markers detected by the 133-plex antibody panel.

**Figure 1.**
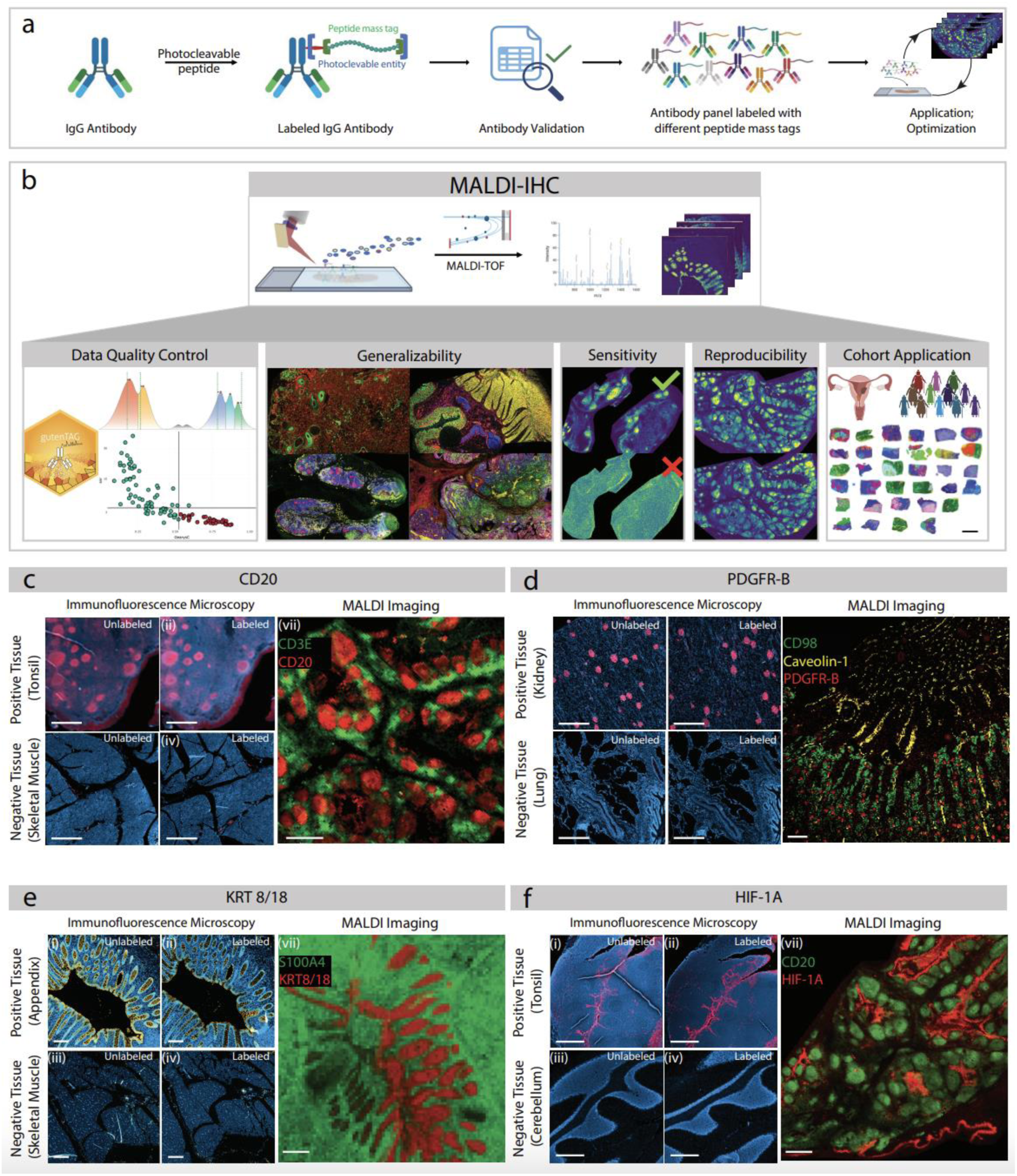
Multiplexed MALDI-IHC: principle, workflow and antibody validation. **a,** MALDI-IHC multiplexing Principle. Each antibody clone is first labeled with a photocleavable mass tag (PCMT) via the amine-N-Hydroxysuccinimide (NHS) click reaction and the specificity of the labeled antibody is confirmed through our validation workflow. Validated antibodies targeting different protein markers are combined into a single staining cocktail and the panel is titrated to identify the concentration that achieves the best SNR in the relevant MALDI imaging conditions. **b,** MALDI-IHC workflow for FFPE samples. 4 µm FFPE tissue sections were mounted on conductive slides), followed by dewaxing and epitope retrieval. After blocking non-specific binding, the full panel of antibodies was applied to the tissue and incubated overnight at 4°C in a humidified chamber. Stained tissues were then thoroughly washed, the slides were dried, and the samples then subjected to UV cleavage to release the mass tags. This was followed by matrix application, rehydration and recrystallization, and then MALDI imaging and data analysis. **(Methods)** **c-f, MALDI-IHC Antibody Validation.** For each antibody, we compared unlabeled and PCMT-labeled versions after immunofluorescence (IF) staining of positive and negative control tissue samples for each marker. This is followed by MALDI imaging on a positive control tissue stained with the PCMT-labeled antibody. An antibody is considered validated only if both the IF and MALDI imaging results match the expected staining patterns, as defined in reference data (e.g., IHC in the Human Protein Atlas or vendor data) and demonstrate a strong signal-to-noise ratio (S/N). Antibody validation examples are shown for CD20 (**c**), PDGFR-B (**d**), KRT8/18 (**e**), and HIF1-A (**f**). In IF, the tested marker is shown in red (**c, d, f**) or yellow (**e**). Positive and negative control tissues used are indicated in each case. MALDI resolution was either 20µm (**e**) or 50 µm (**c, d, f**). Scale bars represent 1 mm (**c, d, f**) or 200 µm (**e**).

We validated the specificity of PCMT-labeled antibodies following a structured workflow (**Fig. 1c-f**) to ensure that PCMT labeling does not alter or destroy an antibody clone specificity. For each target, we compared immunofluorescence (IF) images after staining positive and negative control tissues with both unlabeled and PCMT-labeled versions of a given antibody. In addition, we compared the staining pattern to the known spatial distribution of the targeted antigen in reference samples (e.g., Human protein atlas). Representative examples are shown for the B cell marker CD20 (**Fig. 1c**), the receptor PDGFR-B (**Fig. 1d**), the cytokeratin 8/18 (**Fig. 1e**), and the hypoxia marker HIF-1A (**Fig. 1f**). An antibody was considered validated if it stained the positive control tissue with the expected pattern after PCMT-labeling and showed no staining of the negative control sample. Following successful IF validation, we proceeded to MALDI imaging with the PCMT-labeled antibody on positive control tissues; only antibodies again showing the expected staining pattern and sufficient signal-to-noise ratio (SNR) were considered validated probes. Among all tested antibodies, we found that only one (anti-EGFR) lost specificity upon PCMT labeling, showing that antibodies validated for other imaging modalities have a high likelihood to work for MALDI-IHC.

In an assessment on tonsil, we found that all major immune markers were detectable, including markers for general myeloid cells (CD11b), monocytes (CD14), leukocytes (CD45), B cells (CD20), T cells (CD3, CD4, CD8, CD7), macrophages (CD68, iNOS, CD209, CD163, Arginase 1), NK cells (CD16, CD56, CD7), dendritic cells (CD11c, CD1c), neutrophils (CD66b, MMP9, Lipocalin 2), and the immune checkpoint marker PD-1 (**Fig. S1a**). Low-abundance or rare cell markers such as CD25, MPO, and LAG-3, as well as transcription factors like GATA3 were challenging to detect in MALDI-IHC, as is also the case in IMC without amplification **(Fig. S1b**). In summary, we have generated and validated a 133-plex tumor- and TME-focused, PCMT-labeled antibody panel for use in MALDI-IHC.

### MALDI-IHC data preprocessing and quality control

In order to process and analyze the targeted MALDI-IHC data efficiently, we developed an open-source data pre-processing and analysis pipeline based on Cardinal^27, 32^, called gutenTAG (**Fig. 2a**). The first step in processing MALDI-MSI data involves annotating the peak that corresponds to a specific PCMT. Although each PCMT has a unique and predictable m/z, experimental variability, for example caused by uneven tissue surfaces, can lead to mass shifts and peak broadening. Automated solutions are therefore needed for this process, especially in analysis of large-scale data. GutenTAG uses a Metapeak formation step designed to handle mass shifts and variable peak widths. This process begins with preprocessing spectra, including normalization, smoothing and baseline removal. Automatic peak detection is performed across spectra and aggregated to form a counts histogram of detected peaks, followed by grouping of the detected peak with neighbouring m/z bins into a single Metapeak (**Fig. 2b**). We illustrate the impact of bin width on an image of a kidney tissue stained for caveolin (**Figure S2A**). Visualization using a narrow (±0.100 Da) bin width in the commercial data processing software resulted in the PCMT peak being missed in the lower highlighted area of the image (dashed white rectangle), which is therefore artifactually devoid of signal. Applying a wide bin width (+/− 0.483 Da; middle image) captures the PCMT peak and therefore the signal across the whole image. Notably, this was also achieved in a semi-automated fashion (**Methods**) by gutenTAG’s Metapeak processing.

**Figure 2.**
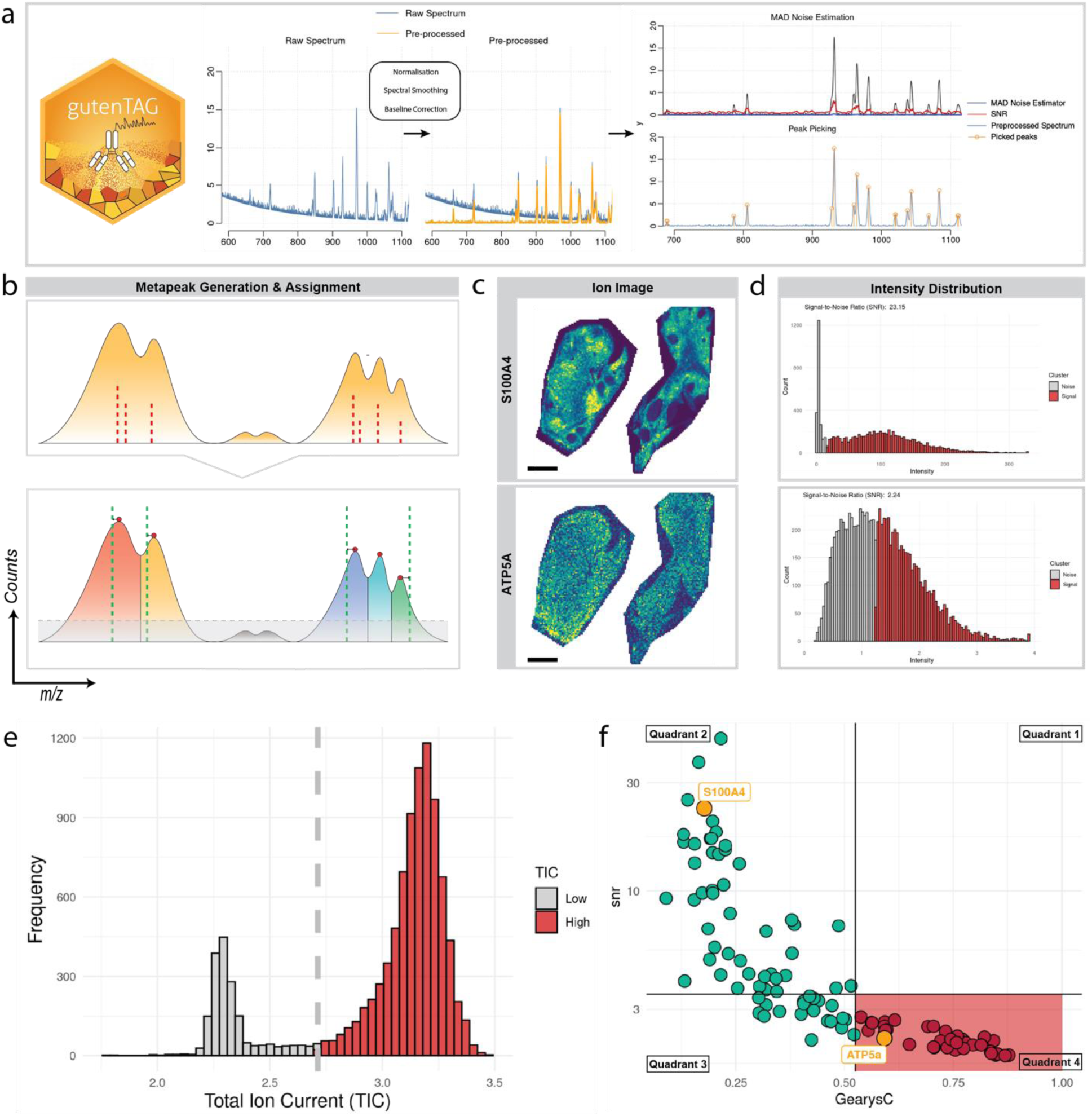
MALDI-IHC data preprocessing using gutenTAG. **a,** Schematic of the initial steps performed by gutenTAG: raw data import, spectral pre-processing including total ion count (TIC) normalization, Gaussian spectral smoothing, local minimum baseline correction, followed by median absolute deviation (MAD)-based peak detection. **b,** Schematic showing Metapeak formation. A Metapeak is generated from the count histogram of detected peaks; gutenTAG groups detected peaks that appear to be in an isotopic envelope into one Metapeak. **c-f,** GutenTAG quality control output for a 50 µm-resolved healthy tonsil MALDI-IHC dataset. **c,** Ion images for S100A4 (top), as an example of a marker with high Signal-to-Noise Ratio (SNR), and ATP5a (bottom), as an example of a marker with low SNR. Scale bar = 1mm. **d**. S/N histogram for the images shown in panel c. The red regions indicate signal and the grey regions indicate noise, as defined using a Gaussian Mixture Model. **e.** Normalized histogram showing the Total Ion Current (TIC) counts for all pixels, where red pixels represent signals, and grey pixels indicate noise. The grey line indicates a threshold determined by Otsu’s method. f. As part of QC, gutenTAG generates the trashcan plot, a decision-making tool showing S/N ratios and Geary’s C values (here with cut-off values of SNR = 3.0, Geary’s C = 0.525). This plot guides users on which staining patterns are as expected; markers in the lower right corner (highlighted in red) in particular should be evaluated as possibly representing poor antibody performance and therefore noise. Markers ATP5a, S100A4, and Ki67 are labeled.

Further, gutenTAG makes semi-automated quantitative assessments of both the overall image and individual ion channels. Thresholding on the total ion current (TIC) histogram distinguishes high-signal from low-signal pixels, thus identifying samples likely to have good signal separation between on- and off-tissue regions (**Fig. 2e**). GutenTAG then calculates the signal-to-noise ratio (S/N) for each marker by applying a Gaussian mixture model (GMM) to the histogram of ion counts to define pixels corresponding to signal and to noise. For example, the marker S100A4 (S/N = 23.15) shows a spatially structured pattern in tonsil, whereas the signal for the marker ATP5a (S/N = 2.24) is seen throughout the tissue (**Figure 2c, d)**. Next, gutenTAG computes Geary’s C autocorrelation scores for each marker. A value much lower or higher than 1 indicates high positive and negative spatial autocorrelation, respectively. Geary’s C values close to 0 indicate high spatial autocorrelation, meaning that the marker has a spatially clustered pattern (e.g., S100A4 in tonsil), whereas values close to 1 indicates no spatial autocorrelation (e.g., ATP5a in tonsil), meaning that the marker has a random or uniform spatial distribution (**Fig. 2f**). We note however that some markers with dispersed spatial patterns, such as Histone H3 and beta-Actin still receive Geary’s C scores close to 0 (0.19 for Histone H3 and 0.21 for beta-Actin) because the score captures local intensity differences (**Fig. S2c, d**). Lastly, gutenTAG summarizes the output from the S/N and Geary’s C analyses to identify markers that are likely to be spatially localized in the images, which informs on whether the marker follows the (expected) staining pattern. For our test tonsil dataset for instance, a S/N cut-off of 3 and a Geary’s C threshold of 0.525 **(Methods)** separated markers into those with high S/N and spatial autocorrelation (quadrant 2, e.g., S100A4), while the fourth quadrant (data points in red) designates those with low values for both these metrics (**Fig. 2f**). The third quadrant includes markers like Ki67, which have low S/N but relatively high spatial autocorrelation, meaning that these may still be reliable signals (**Fig. 2f, Fig. S2b**). In sum, this “trashcan” plot enables evaluation of a marker’s spatial distribution and S/N ratio, supporting further assessment of antibody performance.

Taken together, the gutenTAG package supports the processing and analysis of targeted MALDI-IHC data, including MALDI data preprocessing, PCMT peak detection and evaluation of the performance of individual antibodies.

### MALDI-IHC captures known tissue structures and histological features

To assess the capability of MALDI-IHC to capture known tissue architecture, we applied it to FFPE tissue samples, for which hematoxylin and eosin (H&E)-stained sections were annotated by a pathologist in parallel. For an initial analysis, we applied a 13-plex antibody panel (**Table S4**) to an ovarian carcinoma sample (**Fig. 3a-c**). This panel targeted the ovarian cancer-specific marker mesothelin, six general tumor and epithelial markers (B2M, p53, MUC1, S100A4, and cytokeratin 8/18 and 19), and markers of the PI3K, WNT, and TGF-β pathways (i.e., both active and general forms of CTNNB1, pAKT (Ser473), and SMAD2/3, the innate immune marker Complement C3, and H3K9Ac to indicate transcriptional activation). After pre-processing and QC, we used gutenTAG to spatially segment the MALDI images by clustering pixels according to mass spectrum similarity. For each sample, we determined the appropriate number of clusters by assessing cluster stability across multiple runs and number of clusters (i.e., *k* values) (Methods; **Fig S3**). We adapted the Adjusted Rand Index (ARI) statistic for this purpose, which is typically used to compare two runs of clustering. Here, we ran the k-means clustering 10 times, each time with a different random initialization, computed the pairwise ARI values for each pair of runs and aggregated the results to obtain a mean ARI score. We then chose the number of clusters (k) with the highest ARI score as the most stable number. We identified 4 clusters (**Fig. S3a**) for the ovarian cancer sample; based on the expressed markers, three of these corresponded to tumor and one to stromal tissue, consistent with pathologist annotation of a sequential section (**Fig. 3a, b**). MALDI-IHC image segmentation did not identify pathologist-annotated blood and exudate, due to the absence of the relevant markers in this small panel.

**Figure 3.**
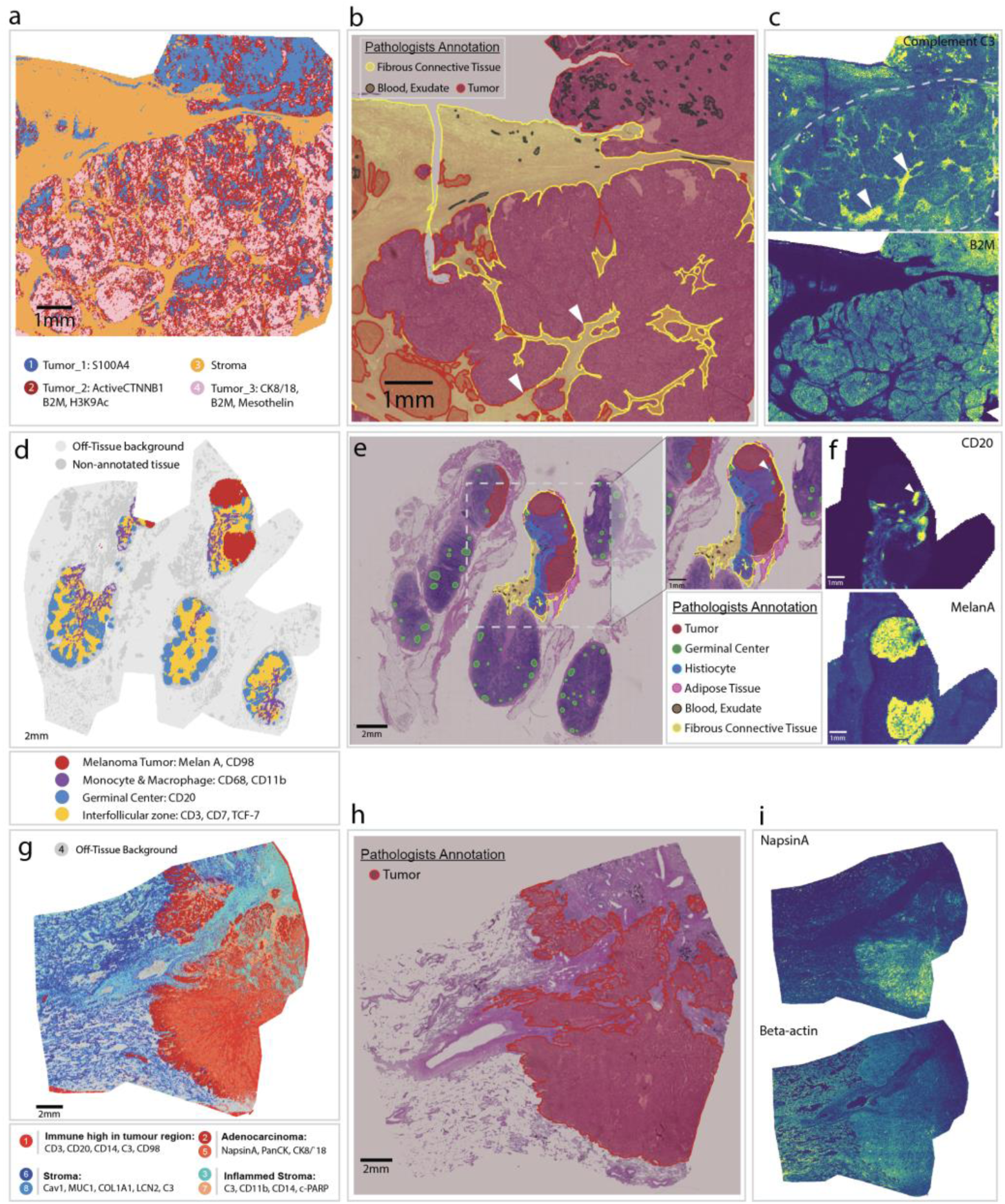
MALDI-IHC captures expected tissue architecture. **(a-c).** 13-plex MALDI-IHC analysis of an ovarian carcinoma sample (20µm resolution). **a**, MALDI-IHC spatial segmentation map indicating the four identified clusters. **b,** Pathologist-annotated H&E image of a consecutive section, in two magnifications. The annotation focuses on the upper right portion of the sample, the lower half including blood and exudate regions is not fully outlined. White arrows indicate two fibrous connective tissue areas. **c**, Ion images for the indicated markers. The grey dashed line in the upper panel of (**c**) outlines the tumor region. **(d-f)**. 120-plex MALDI-IHC analysis of a lymph node melanoma metastasis (50 µm resolution). **d,** MALDI-IHC spatial segmentation map, indicating the twelve identified clusters, highlighting the melanoma tumour, germinal center, intergerminal region and macrophage/monocyte-rich clusters. Remaining clusters corresponding to on-tissue pixels are labelled non-annotated and coloured in dark grey, while off-tissue background clusters are coloured in light grey. **e,** Pathologist-annotated H&E image of a second section of the same sample, in two magnifications. Detailed annotations are shown only for the central portion of the tissue (shown also in higher magnification). The white arrow points to the germinal centre. **f**, Ion images for the indicated markers. (**g-i**) 120-plex MALDI-IHC analysis of a lung adenocarcinoma sample (50µm resolution). **g,** MALDI-IHC spatial segmentation map indicating the eight identified clusters. **h,** Pathologist-annotated H&E image of a separate section, showing tumor regions in red. **i**, Ion images for the indicated markers.

In a second analysis, we used a 120-plex tumor panel (**Table S2**) to stain a melanoma lymph node metastasis sample (**Fig. 3d-f**) and lung adenocarcinoma (LUAD) tissue (**Fig. 3g-i**). We identified twelve clusters in the melanoma sample (**Fig. S3b**), which we annotated as tumor, lymph node components including the germinal centre, interfollicular zone, a macrophage and monocyte-rich region, and several clusters corresponding to off-tissue regions (**Fig. 3d**). Pathologist annotations again showed strong agreement with the regions identified in our MALDI-IHC segmentation (**Fig. 3e**), in particular the tumor, connective tissue, germinal centre, and interfollicular zone, despite the H&E data originating from a distant tissue section. CD20 staining closely aligned with the germinal centre cluster structure and Melan A was prominent in the melanoma tumor cluster, as expected (**Fig. 3f**). The pathologist-annotated histiocyte-rich region (**Fig. 3e**, blue area), corresponded to a macrophage and monocyte-rich cluster in our MALDI-IHC segmentation, characterized by high expression of CD11b and CD68 and therefore aligning well with a histiocyte marker profile. Pathologist-annotated blood/exudate regions were absent in our MALDI analysis, likely due to the spatial separation of the samples. In the lung adenocarcinoma (LUAD) sample, we identified eight clusters, including tumor, an immune-rich cluster within the tumor, inflamed stroma, healthy lung components, and an off-tissue cluster (**Fig. 3g-i; Fig S3c**). LUAD-specific markers such as Napsin A were again strongly detected in the tumor clusters, which also aligned closely with pathologist annotation; minor discrepancies were expected since the images were not derived from a serial section of the sample.

Overall, our findings demonstrate that spatial segmentation of MALDI-IHC data identifies regions that correspond to pathologist-annotated histological images, indicating that MALDI-IHC captures expected tissue architecture.

### Reproducibility assessment of whole-slide MALDI-IHC imaging

Next, we evaluated the intra- and inter-slide reproducibility of MALDI-IHC imaging. We analyzed consecutive sections of tonsil (n=2, 97 mm² each) on a single slide, colon (n=6, 37 mm² each) distributed across two slides with three sections per slide, and placenta (n=2, 180 mm² each) on two slides (**Fig. 4a**). Reproducibility was assessed by comparing replicate samples in terms of: (1) identified markers, (2) marker intensity, (3) spatial pattern, and (4) cluster composition.

**Figure 4.**
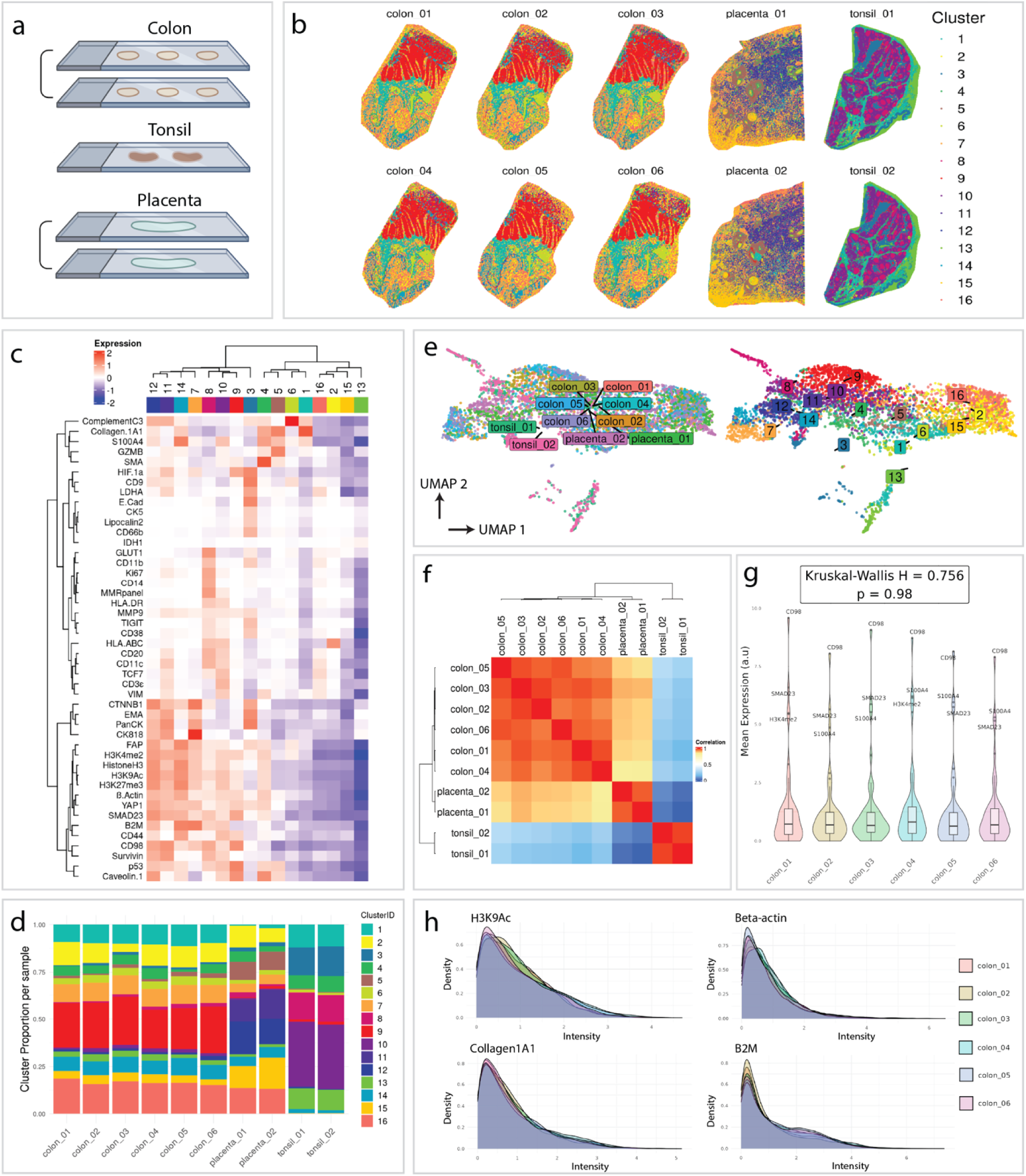
Reproducibility assessment of MALDI-IHC. **a,** The schematic shows three groups of samples used to assess reproducibility of MALDI-IHC. Depicted are serial sections of colon (n=6) on the same or different slides (top), serial sections of tonsil (n=2) on the same slide (middle), and serial sections of placenta (n=2) on different slides (bottom). **b,** MALDI-IHC analysis of samples depicted in a and stained with a 120-plex panel. Clusters were identified on data from all ten samples. Depicted are the sixteen identified clusters mapped to each sample, with each cluster indicated by color. **c,** Heatmap illustrating marker expression in each identified cluster. Euclidean distance was used as the distance measure for rows (markers) and ward.D2 was used to cluster columns (clusters). **d,** Stacked bar plot showing the proportion of each cluster per sample. Colours are as in b. **e,** UMAP representations of the MALDI-IHC dataset with sample ID and cluster number indicated. **f,** The heatmap shows Pearson correlation between samples based on their marker expression profiles; correlation coefficients range from 0 (blue) to 1 (red). The dendrogram shows hierarchical clustering with a complete linkage approach. **g**, Violin plots show mean expression of all markers across the 6 colon replicates. There is no significant difference between replicates. **h**, Signal intensity/density plots for selected markers across the entire imaging area of the colon samples, with each sample indicated by color.

All samples were stained in a single batch using a 120-plex MALDI-IHC panel, and then also imaged in a single batch; 46 of the markers were relevant for these tissues (**Table S5**). gutenTAG quality control was performed identically on the data from each sample, followed by integration into a single dataset encompassing all samples. We used k-means clustering of this dataset to segment the images based on pixel similarity, choosing to define 16 clusters based on our cluster stability metric (**Fig. S3d**); clusters 3, 8, 10 and 13 were unique to tonsil and clusters 2, 7, 9, 11, 12, 14, 15 and 16 were found in colon and placenta (**Fig 4b, d**). The clusters corresponded with expected histological features of each tissue. For example, in tonsil, both spatial and marker patterns are consistent with cluster 10 representing non-germinal center regions, characterized by high expression of CD3, CD44, and TCF7, and cluster 8 representing germinal centers, marked by elevated CD20, CD11b, and HLA-DR signals (**Fig. 4b, c**). In colon and placenta, clusters 7 and 11 exhibited epithelial features (CK8/18, PanCK), while clusters 4 and 5 were associated with submucosa (colon) and decidua (placenta), displaying markers such as Collagen-1A1 and SMA. Cluster 12 showed mucosal characteristics, with markers such as B2M, S100A4, CTNNB1, and SMAD2/3. Reassuringly, replicates for each tissue type broadly showed the same clusters and spatial distribution (**Fig. 4b**), mapped near each other in the UMAP (**Fig 4e**), comprised very similar cluster proportions (**Fig. 4d**), and were well correlated in terms of marker composition (**Fig 4f**).

All markers for a given tissue were consistently identified across replicates, and there was no significant difference in average intensity for any of the three tissues (**Figure 4g; Figure S4a, b**). To obtain a better overview of the ion image for each marker, we plotted the signal intensity of each marker against its density across the imaged area and compared the resulting histograms across replicates. This revealed consistent intensity distributions across replicates for most markers (**Fig. 4h, Fig. S4c**), further supporting the reproducibility of MALDI-IHC. We note however that not all curves fully overlapped, with discrepancies observed both within and between slides (**Fig S4c**, see markers CD98 and Caveolin1). In conclusion, although a few markers show suboptimal quantitative patterns, these analyses largely show reproducibility in MALDI-IHC, with reliable marker detection, ion image generation, and cluster identification across replicates.

### 120-plex MALDI-IHC analysis of an endometrial cancer cohort

Having established our experimental and data analysis pipeline, we next applied 120-plex MALDI-IHC to a patient cohort comprised of 18 patients with endometrial cancer (EC) (**Table 2**). EC is the most common gynaecological malignancy in developed nations, with a rising incidence rate. EC can be stratified into four molecular subtypes based on genomic information^33,43^: POLE-ultramutated, microsatellite instability-high (MSI), copy-number low (CN-Low), and copy-number high (CN-High). POLE-ultramutated tumors show the most favourable prognosis and CN-High tumors, frequently harbouring TP53 mutations, are associated with the poorest outcomes. Despite substantial progress in understanding the genomics and transcriptomics of EC^44–45^, its spatial proteomic landscape remains underexplored, with only a limited number of bulk proteomics studies conducted to date^46–47^. As a first step towards this goal, we employed our 120-plex MALDI-IHC panel for highly multiplex protein imaging of a small EC cohort including all TCGA subtypes.

**Table 2.** Clinical features of the endometrial cancer cohort.

|  | POLE | MSI | CN-High | CN-Low |
| --- | --- | --- | --- | --- |
| Patient Count | 4 | 8 | 2 | 4 |
| Sample Count | 9 | 12 | 3 | 5 |
| Average Age | 53.0±1.4 | 69.6±14.5 | 76.7±6.7 | 57.8±14.4 |
| Matched Healthy Control | 1 | 5 | 0 | 0 |
| Average BMI | 26.5±3.1 | 31.3±1.8 | 31.0±0.0 | 29.5±4.0 |
| Stage 1-2 Patient | 4/4 | 4/8 | 2/2 | 4/4 |
| Grade 1-2 Patient | 1/4 | 7/8 | 0/2 | 4/4 |
| Metastasis and/or Lymphovascular invasion | 1/4 | 4/8 | 0/2 | 1/4 |

We analysed 33 formalin-fixed, paraffin-embedded (FFPE) samples from 18 patients, comprising all four molecular subtypes (**Table 2**). These included 1-3 tumor samples per patient along with matched healthy control tissue from a subset of patients (4 MSI, 1 POLE). We performed the MALDI-IHC workflow for this cohort with whole-slide imaging of 120-plex stained samples at 50 µm using a RapifleX MALDI-TOF tissue typer and generated a dataset of 4,320,442 pixels and covering a total area of 10,801.1 mm² (**Fig. 5a**). Following preprocessing and marker panel refinement with gutenTAG, we reduced the panel to the 70 most informative markers (**Table S6**), mainly by removing low quality channels flagged by gutenTAG’s S/N score and Geary’s C criteria. We also removed 9 samples that showed low staining intensity across all markers; the remaining 24 samples corresponded to 16 patients.

**Figure 5.**
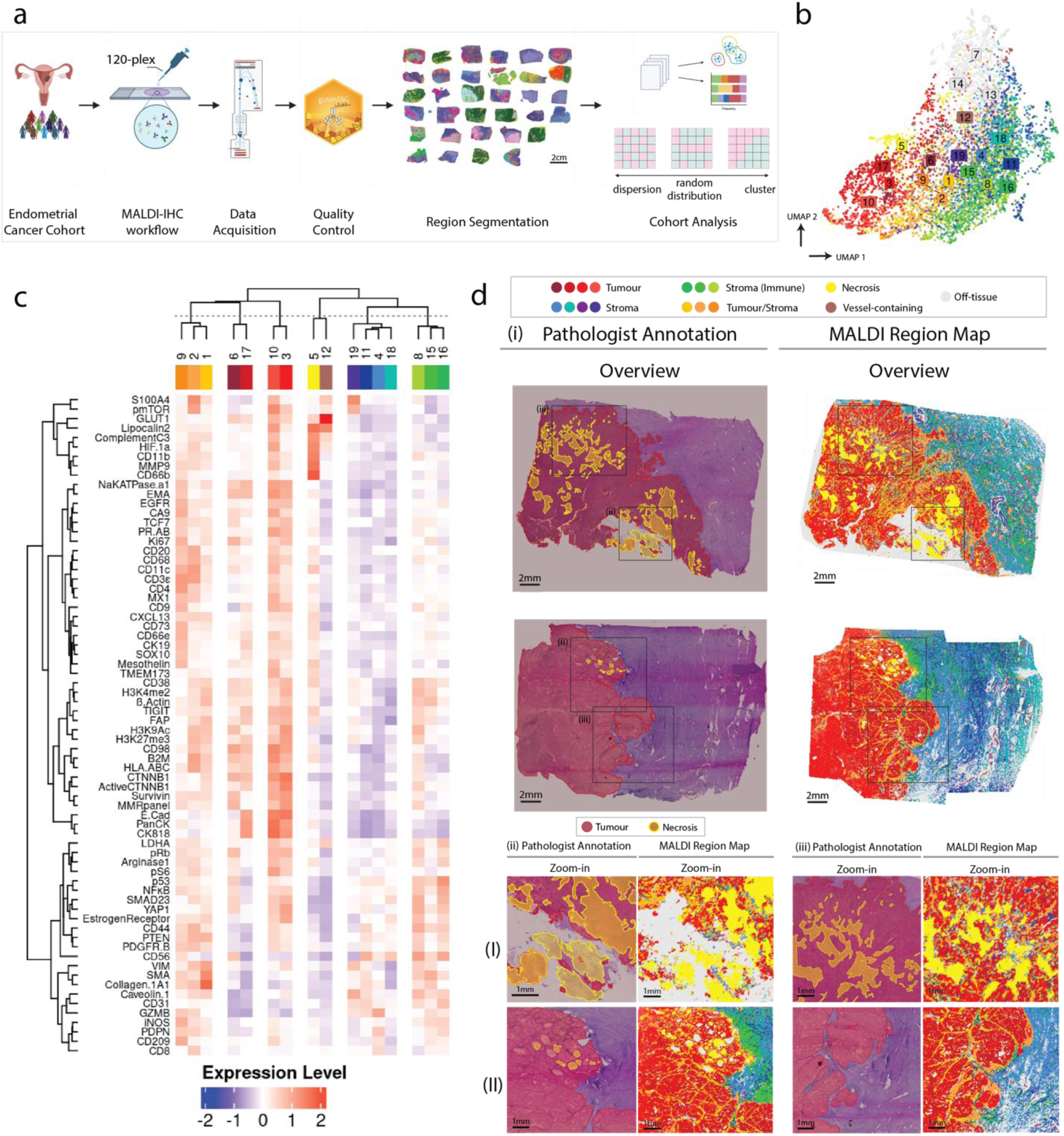
High-plex MALDI-IHC analysis of an endometrial cancer cohort. **a,** Overview of the MALDI-IHC experimental and data analysis workflow for an endometrial cancer cohort. Samples were processed in three separate batches, each stained with a freshly prepared cocktail of 120 MALDI-IHC antibodies. Whole-slide imaging was then performed at a 50µm spatial resolution. Preprocessing and quality control of the raw data were conducted using the gutenTAG pipeline, and high-quality markers were subsequently used to identify pixels with similar mass spectra, which were spatially mapped onto the samples. The generated clusters were then analyzed in terms of marker expression and spatial autocorrelation. **b,** UMAP projection of the 70 markers colored by identified clusters. **c,** The heatmap shows z-score normalized marker intensities for all nineteen identified clusters. Clusters annotated as off-tissue were excluded from the analysis. Hierarchical clustering was applied to both rows (markers) and columns (clusters) using Euclidean distance for rows and ward.D2 for columns. **d,** Validation of segmented regions with pathologist annotations. Two representative samples are shown. A pathologist-annotated H&E image and the corresponding MALDI cluster map are shown pairwise for the full image (i) and for two zoomed in views (ii, iii). For sample I, the H&E image is of a consecutive section to that imaged by MALDI-IHC; for sample II, the H&E and MALDI-IHC images are on the same section. Pathologist annotations were conducted independently, outlining the tumor (red) and necrotic (yellow) regions; the latter can include some areas where mucus, hemorrhage, exudates and necrotic material are intermixed. The 19 annotated clusters are indicated by colour.

We then applied our correlation distance k-means clustering (Fig S3e) to reveal 19 clusters based on mass spectral marker profiles. We annotated these clusters based on their marker patterns as stromal (n=7), 3 of which had higher immune content (stroma/immune), tumor (n=4), mixed stromal-tumor (n=3), off-tissue (n=3), vessel-containing (n=1) and necrotic (n=1) (**Fig 5b, c**). Clusters in these different annotated classes were separated on a UMAP (**Fig. 5b**). Tumor clusters 3, 6, 10 and 17 exhibited high expression levels of several tumor-associated markers, including epithelial markers such as PanCK, E-Cadherin, and CK8/18, as well as elevated expression of cell state markers like Ki67 (indicative of cell proliferation) and Survivin (inhibition of apoptosis), metabolism-related markers like GLUT1 (glucose transport), and pathway markers such as CTNNB1, representing WNT signalling (both full-length and activated forms). Stromal clusters expressed high levels of stromal markers such as PTEN (often mutated in tumors), p53, caveolin, and collagen-1A1. Three of these stromal clusters (8, 15, 16) showed higher expression of these stromal markers, along with immune markers like iNOS (associated with macrophages) and NFKB, pathway markers such as YAP1 (Hippo signalling) and the cancer stem cell marker CD44. In contrast, stromal clusters 4, 11, 18 and 19 exhibited lower levels of lymphocyte markers such as CD20, CD3ε, CD4 and TCF7. Interestingly, stromal cluster 19 exhibited high expression levels of phospho-mTOR and S100A4, indicating a tumor-like characteristic. Clusters 1, 2 and 9 strongly expressed tumor, stromal and immune markers and were found largely within the tumor area or on tumor borders and were therefore annotated as tumour/stroma clusters.

We spatially mapped our annotated clusters onto the samples and compared their patterns to independent pathologist annotations of an H&E image of the same or a sequential section (**Fig 5d**). Importantly, major tissue features, namely the tumor mass, necrotic regions, and stromal areas, showed strong concordance between MALDI-IHC-identified and pathologist annotated regions. For example, pathologist-annotated necrotic tissue (yellow-shaded regions) aligned well with the MALDI-IHC segmentations (cluster 5, yellow) (**Fig. 5d**). Some discrepancies are expected as areas with necrotic regions may include mucus, haemorrhage and exudates, and precise boundaries are challenging to demarcate. The MALDI-IHC tumor regions mapped broadly onto the pathologist’s manual annotations but had greater granularity due to the molecular information included. We note that the samples differed in their tumor content, with some samples composed largely of healthy endometrial tissue (**Fig. S5a, b**). Most of the tumor, stromal and tumor-stromal clusters were present across all samples, with tumor regions absent in healthy samples as expected (**Fig. S6**). An analysis of global spatial autocorrelation using the Moran’s I statistic **(Methods)** showed that cluster 2 (tumor-stromal), 6 (tumor), 5 (necrosis) and 12 (vessel-containing) were more spatially structured in the most aggressive CNHigh subtype, whereas this was true for cluster 10 (tumor) in CNLow (**Fig. 6, S6**). However, given the small size of this cohort, these results must be explored in larger studies. Notably, spatial autocorrelation across samples and clusters varied widely, reflecting spatial heterogeneity even within this small cohort.

**Figure 6.**
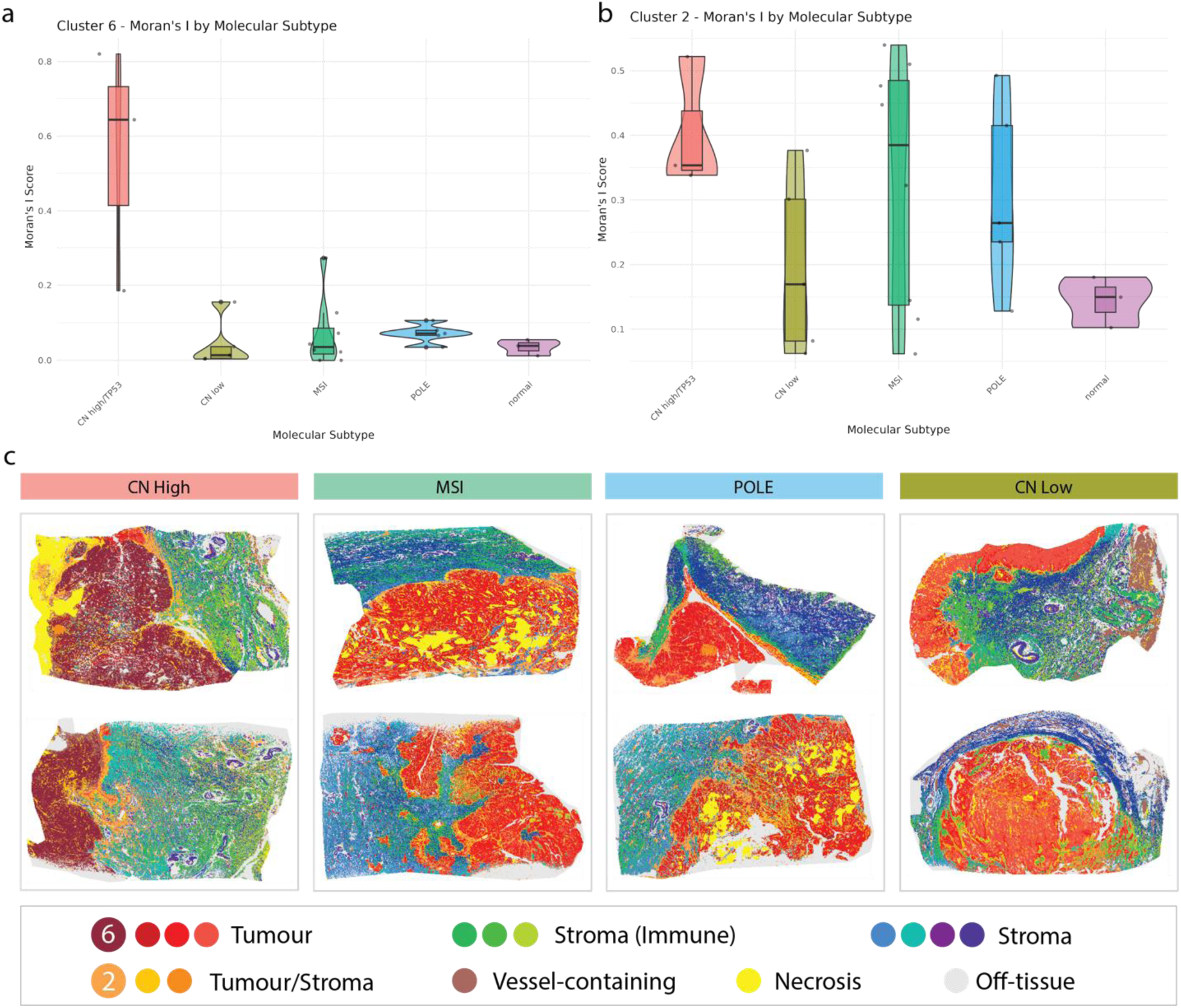
Spatial analysis of segmented regions across endometrial cancer molecular subtypes. **a, b,** Violin plots depicting the distribution of Moran’s I statistic for the tumor cluster 6 (**a**) and the tumor-stromal cluster 2 (**b**) across molecular subtypes. **c,** Representative spatial patterns of the identified clusters are shown across the four molecular subtypes, indicated by colour.

Our data demonstrate that 100+ plex MALDI-IHC can be applied to samples from a patient cohort and can identify expected histopathological features with added molecular granularity.

## Discussion

Highly multiplexed proteomics imaging has become a cornerstone of spatial biology, offering direct protein evidence that complements spatial transcriptomics data. Traditionally, the multiplexity of these methods (e.g., IMC) in routine application has been around 40-plex, but this often falls short in capturing the full complexity of cellular phenotypes, states, and relevant biological pathways within intricate systems. While cyclic-staining approaches using immunofluorescence can enhance multiplexity, they are hindered by lengthy procedures and the risk of specimen damage and deformation after multiple staining cycles. And although sequencing-based methods also enable high-plex protein marker detection, they are time- and resource-intensive. In this work, we demonstrate the potential of MALDI-IHC to achieve simultaneous imaging of over 100 protein markers, in a rapid, efficient and scalable manner.

To achieve this, we began by labeling antibodies previously used in our IMC studies, which had shown robust staining performance and combined them with available pre-conjugated PCMT **(Methods)**. We established an antibody validation workflow to ensure the specificity and sensitivity of MALDI-IHC antibodies. Notably, almost all antibodies demonstrated high specificity, indicating that the MALDI-IHC PCMT labeling process is a mild modification that does not compromise antigen recognition. The high success rate in antibody labeling suggests that future bottlenecks in expanding MALDI-IHC multiplexity will likely lie in the availability of highly sensitive and specific unlabeled IgG antibodies and the development of additional PCMTs. Furthermore, the use of stable isotopically pure PCMTs could theoretically triple the number of available tags, as each PCMT would appear at a single m/z location, leaving the three isotopic peaks location accessible for other PCMTs. We also report an optimized sample preparation workflow in which we reduce the intensity of sodium adduct peaks from high-intensity PCMT peaks, which complicate data analysis and limit practical multiplexity, by vigorous washing with volatile salt buffer.

To address data analysis challenges of MALDI-IHC data, we developed gutenTAG, an R-based open-source data pre-processing, quality control, and analysis pipeline. gutenTAG groups together *m/z* bins into a single Metapeak based on a watershed segmentation algorithm. This groups all isotopic peaks from a PCMT into a single metapeak, effectively performing deisotoping. This approach is well-suited to MALDI-IHC, as the protocol introduces minimal interfering peaks, allowing for robust Metapeak formation. Additionally, the broader Metapeak width semi-automatically resolves problems with incomplete representation of ion images that can occur with too-narrow bin widths. gutenTAG further implements quantitative methods for quality control of each ion channel image, employing a “trashcan” plot that combines S/N separation and Geary’s C score as a guide to assess antibody staining patterns vis-à-vis the expected distribution for that marker. This renders the traditionally manual QC process more reliable and less labor-intensive.

Since we performed MALDI imaging at supracellular resolutions (20-50µm), downstream analyses were done at the pixel/region level. We identified pixels with similar mass spectral profiles by k-means clustering using a correlation distance metric on gutenTAG processed data, which could then be spatially mapped to the analysed samples. We found that the correlation clustering metric emphasizes the overall similarity of the spectrum and is more robust to intensity differences across acquisitions than Euclidean distance. This implementation allowed us to cluster over 4 million pixels from the endometrial cancer cohort in ∼1 hour and identify shared biological features across samples. As such, it therefore presents a flexible, reproducible and scalable region segmentation algorithm for MALDI-MSI data. The open-source nature of our code also facilitates ongoing optimization and the integration of additional analysis modules.

With a foundational framework established for high-plex MALDI-IHC imaging, we applied it across a range of tissue types (ovarian cancer, melanoma lymph node metastasis, lung adenocarcinoma, endometrial cancer), confirming that MALDI-IHC is a robust method adaptable to various FFPE tissue sections without experimental modifications. This highlights a key advantage of MALDI-IHC over other high-plex IF methods, which often necessitate extensive optimization for each panel or tissue type, especially regarding markers’ staining orders in cyclic approaches. MALDI-IHC reliably identified histopathological features that closely matched pathologist annotations, further validating its reliability. For instance, tumor regions in all tested samples were accurately detected, alongside the identification of lymph node structures in melanoma samples and adenocarcinoma regions in the LUAD sample. All identified regions displayed expected protein marker expression profiles. Notably, in the melanoma sample, MALDI-IHC identified germinal center structures that were not visible to the pathologist in H&E-stained sections but appeared correctly annotated based on CD20 and Ki67 staining; this is most probably because the structures of germinal centers were compromised in these samples due to tumor infiltration. Furthermore, in the ovarian cancer and LUAD samples, MALDI-IHC successfully segmented sub-regions within tumor and stroma based on cell state and immune profiles – features challenging to capture through conventional H&E or low-plex approaches.

We further evaluated the reproducibility of MALDI-IHC to assess its suitability for cohort studies. We observed excellent intra- and inter-slide qualitative reproducibility but noted some variability in inter-slide intensities for a few markers. Factors such as matrix coating differences and variations introduced during the manual matrix rehydration-recrystallization step may contribute to this variability. Future improvements should focus on developing a more controlled and automated matrix rehydration-recrystallization process. Additionally, introducing control samples adjacent to the samples of interest and performing matrix spraying with a series of mass tag standards mixed with the MALDI matrix could help correct for slide-to-slide signal intensity differences.

Finally, we applied high-plex MALDI-IHC imaging on a clinical cohort of 33 endometrial cancer samples, acquiring whole-slide imaging data for all samples at 120-plex in one week of active measurement time. On the data analysis front, we confirmed that gutenTAG efficiently manages large datasets, performing pre-processing, quality control and region segmentation based on k-means clustering using this tool. We could identify tumor, stromal and mixed clusters and showed in initial exploratory analyses that spatial structure of some clusters differed across endometrial cancer molecular subtypes. We have thus demonstrated the feasibility of performing 100+ plex whole slide imaging in a scalable manner, indicating that the MALDI-IHC method may be effectively applied to larger cohorts in the future.

In summary, we have established and validated 100+plex MALDI-IHC imaging and have shown that it can be applied for cohort studies. The substantial richness of the data, even when applying the same high-plex panel to different tissues or disease types, underscores the potential of >100-plex MALDI-IHC technology. The method holds significant promise for uncovering novel tissue regional phenotypes and elucidating region-region interactions. Future studies with a more targeted MALDI-IHC panel and larger sample sizes will offer deeper biological insights into the systems under investigation. Additionally, with ongoing technical advancements, we expect MALDI-IHC to evolve towards achieving sub-micron spatial resolution, which would expand its capabilities from regional and spatial pattern analysis to providing detailed cellular-level insights. This evolution will further enhance our understanding of complex biological systems and drive forward the discovery of new biomarkers and therapeutic targets.

## Methods

### Human tissue samples collection

Formalin-fixed, paraffin-embedded (FFPE) human tissue blocks for method establishment, optimization and validation experiments were obtained from the Gewebe-Biobank at the Institute of Pathology and Molecular Pathology (IPMP), Zürich. All tissue samples are anonymized and were collected with informed consent in compliance with ethical standards and regulatory guidelines for human research.

Formalin-fixed, paraffin-embedded (FFPE) human endometrial cancer tissue blocks for the cohort study were obtained from the archive of the Institute of Pathology from the University Hospital Basel. Each sample is linked with clinical information provided solely for research purposes, under ethical approval and in compliance with regulatory guidelines for human research. All samples were collected with informed consent, and patient data has been anonymized to protect privacy. The study was approved by the Ethics Commission of Northern Switzerland (EKNZ; study ID: 2024-01079).

### PCMT-Antibody Preparation

#### Antibodies collection

All antibodies used in this study are summarized in the **Supplementary Table 1**.

#### Reagents and Buffers

Single-use PCMT aliquots, containing a dried mass-tagging reagent, were obtained from AmberGen Inc. and were stored at ≤ −20°C in desiccated, light-protected conditions and equilibrated to room temperature prior to use. Anhydrous N,N-Dimethylformamide (DMF) was used to reconstitute the reagent immediately before antibody labeling, and PBS (pH 7.5) served as the diluent. Additional buffers included 1 M sodium bicarbonate, 1 M glycine, and Tris-buffered saline (TBS, 50 mM Tris, 200 mM NaCl, pH 7.5), which were all prepared in-house and dissolved in LC-MS grade water (Thermo Scientific Chemicals, ALF047146). The antibodies were first purified if they contained protein carriers or stabilizers from the suppliers.

#### Antibody Purification

Antibodies (100 µg) were purified using a 50 kDa MWCO Amicon Ultra-0.5 Spin filter unit (Millipore-Sigma, UFC5050) with PBS as the wash buffer. Each purification cycle involved adding 450 µL of PBS to the antibody solution, followed by centrifugation at 12,000g for 10 minutes. This wash step was repeated three times to ensure thorough purification. The final antibody was recovered and resuspended in PBS to the concentration of 1 µg/µL.

#### Labeling Procedure

Labelling antibodies with PCMTs follow the manufacturer’s instructions (AmberGen Inc.). Purified antibodies (100 µg) were mixed with 1 M sodium bicarbonate (100 mM final) and then immediately added to reconstituted PC-MT aliquots, incubating for 1 hour with gentle mixing, protected from light. The reaction was quenched with 1 M glycine (100 mM final) for 15 minutes. Unreacted reagent was removed using a 50 kDa MWCO Amicon Ultra-0.5 spin filter unit, with six buffer-exchange cycles of 450 µL TBS, each at 12,000g for 10 minutes.

#### Post-Labeling Preservation

The labeled antibody solution was mixed with glycerol (40% final), sodium azide (0.05% final), and protease inhibitors as well as EDTA stock (both from Halt™ Protease Inhibitor Cocktail, Thermo Fisher, 1X final), yielding a final concentration antibody at 250µg/mL in TBS. The final products, termed PCMT-antibodies, were stored at −20°C in light-protected containers.

### FFPE tissue section preparation

FFPE blocks were sectioned at a thickness of 4 µm using a rotary microtome. The sections were mounted onto IntelliSlides (Bruker) coated with 0.01% Poly-L-Lysine, then baked overnight at 37°C, followed by an additional 2 hours at 60°C to enhance tissue adherence. For storage, the sections were kept at 4°C for short-term or at −20°C for long-term preservation.

### MALDI-IHC sample preparation

Tissue sections were equilibrated to room temperature for approximately 1 hour in a desiccator chamber before deparaffinization. Deparaffinization was performed on an AS-2 carousel (Pathisto) with the following program: three 10-minute cycles in UltraClear™ (J.T.Baker), followed by two 5-minute washes in 100% ethanol, two 3-minute washes in 96% ethanol, two 3-minute washes in 90% ethanol, one 3-minute wash in 80% ethanol, and one 3-minute wash in 70% ethanol. The slides were then soaked in TBS buffer for at least 5 minutes. Deparaffinized slides were subjected to heat-induced antigen retrieval (HIER) in a basic buffer (10 mM Trizma, 1 mM EDTA, pH 9.2) and heated at 95°C for 30 minutes in a decloaking chamber (BioCare Medical). Slides were then cooled at room temperature for 20 minutes, washed in TBS for 15 minutes, and regions of interest were outlined using a hydrophobic pen (Vector Laboratories). Samples were blocked for 1 hour at room temperature in a humidified chamber with a MALDI-IHC-specific blocking buffer containing 5% (v/v) normal horse serum (Gibco), 5% (w/v) BSA, and 0.05% (w/v) Octyl β-D-glucopyranoside (OBG) detergent in TBS.

The antibody solution was prepared in blocking buffer and diluted to the optimal staining concentration, as determined by titration studies, typically between 0.5 µg/mL and 2 µg/mL. All antibodies in the staining panel were combined into a single staining cocktail, which was applied to the tissue region within the hydrophobic barrier. Tissues were incubated with the staining cocktail at 4°C for a minimum of 16 hours. Excess antibodies were washed off with three 5-minute washes in TBS (prepared in LC-MS grade water) by vigorously shaking the Coplin jar (sealed and shaken at 300 rpm on an Eppendorf Thermomixer), followed by two 3-minute washes in 50 mM ammonium bicarbonate buffer (prepared in LC-MS grade water). Finally, slides were briefly dipped in LC-MS grade water for 10 seconds before being dried in a UV-blocking vacuum desiccator chamber (Cleatech, 1300-2-B) at –0.08 MPa for at least 2 hours. The dried slides were then photocleaved using a UV exposure (Modell II, 2 × 6 W, Roth AG) and exposed at 355nm for 10 minutes.

To prepare the MALDI matrix, α-Cyano-4-hydroxycinnamic acid (CHCA) was dissolved in a solution of 70% acetonitrile (ACN), 0.1% trifluoroacetic acid (TFA), and 10 mM ammonium phosphate in LC-MS grade water. Additionally, angiotensin II (ProteoMass™, Sigma-Aldrich) was added to the CHCA matrix solution at a final concentration of 20 nM as an internal standard. A TM5 matrix sprayer (HTX Technologies) was used to spray the photocleaved slides with the following spraying parameters: tray temperature at 25°C, nozzle temperature at 60°C, flow rate of 0.1 mL/min, nozzle velocity at 1350 mm/min, eight passes, track spacing of 3 mm, nozzle height 40 mm above the slide, “CC” movement pattern, nitrogen gas pressure at 10 psi, and gas flow rate at 2 L/min. A 10-second drying time was applied between each pass.

The sprayed slides underwent rehydration-recrystallization by affixing the CHCA-coated slide to the bottom of a 100 mm ′ 20 mm plastic petri dish (Corning, #430167) and placing a filter paper soaked with 1 mL of 5% isopropanol (IPA) solution on the lid. The petri dish was inverted, positioning the filter paper at the bottom with the coated slide facing downward toward the filter paper. This self-made humid chamber was placed in a 55°C oven, and the slide was rehydrated for precisely 105 seconds. Immediately afterward, the slide was transferred to a vacuum desiccator and dried under vacuum (−0.08 MPa) for at least 5 minutes.

### MALDI-IHC panel titration

The 120-plex antibody panel was combined into one staining cocktail for titration. Antibody titrations were conducted at concentrations of 0.1, 0.5, 1, and 2 µg/mL on serial sections from various tissue types, including tonsil, kidney, colon, and melanoma. For 28 markers that presented low abundance or were difficult to detect via MALDI-IHC (**Table S3**), the titration range was extended to 5 µg/mL. All MALDI-IHC data were acquired at a spatial resolution of 50 µm, and for each marker, the maximum signal-to-noise (S/N) ratio was assessed across titration concentrations and tissue types to identify optimal detection conditions.

### MALDI-TOF imaging

CHCA matrix-coated slides were spotted with 1 µL of a red phosphorus-acetone mixture and subjected to MALDI-TOF imaging. Imaging was performed on a rapifleX MALDI-TOF-MS instrument (Bruker Daltonics) with the following parameters: reflector mode, positive polarity, laser spot sizes of 20 or 50 µm with continuous raster scanning at corresponding resolutions, 300 laser shots per pixel, approximately 70% laser power, and normalization to total ion count (TIC) for generated images. Image and spectral analysis were conducted using gutenTAG and SCiLS Lab software (Bruker Daltonics).

### Post-MALDI H&E staining

After MALDI-TOF imaging, slides were subjected to matrix removal by washing in 70% ethanol for 3 minutes, repeated three times. Slides were then rehydrated in TBS for at least 10 minutes before staining. Hematoxylin and Eosin (H&E) staining was performed using a Thermac CellStain-6 auto-stainer with the following program: 0.1% hematoxylin (Mayer’s Hematoxylin, Dako) for 3 minutes, followed by a 1-minute rinse in distilled water (ddH₂O). Slides were then dipped in a differentiator solution (70% ethanol, 0.3% hydrochloric acid) for 1 minute, rinsed in ddH₂O for 1 minute, followed by 1 minute in Scott’s tap water (0.2% sodium bicarbonate and 2% magnesium sulfate, both in w/v), and another rinse in ddH₂O for 1 minute. This was followed by a 1-minute dip in 95% ethanol, staining with 1% (w/v) Eosin Y (Apollo Scientific) for 1 minute, and a 1-minute rinse in Scott’s tap water. Slides were then sequentially dehydrated in 95% ethanol for 1 minute and 100% ethanol for 2 minutes, followed by clearing in UltraClear™ for 4 minutes. Coverslips were mounted with Eukitt (Sigma Aldrich) and allowed to harden. Whole-slide imaging (WSI) was performed on a Zeiss Axioscan Z1 slide scanner under bright-field illumination at 10x magnification to acquire H&E images.

### Immunofluorescence Microscopy antibody validation

Samples were prepared as described for MALDI-IHC through the antigen retrieval step. For antigen blocking, samples were incubated in PBS containing 3% (w/v) BSA (Sigma) and 0.1% (v/v) Triton X-100 (Sigma) for 1 hour at room temperature in a humidified chamber. Single-plex antibody staining solutions were prepared by diluting PCMT-labeled or unlabeled antibodies in blocking buffer to a concentration of 2 µg/mL, or as per the vendor’s recommendations. Tissue sections were incubated with the antibody staining solution at 4°C overnight. Excess antibody was removed by washing slides in a Coplin jar with TBS for 3 cycles of 5 minutes each. A secondary antibody, specific to the host species of the primary antibody (all obtained from Invitrogen’s Alexa Fluor series), was then applied at a 1:500 dilution in TBS and incubated at room temperature for 1 hour. Following incubation, the secondary antibody was washed off with three 5-minute washes in TBS. Nuclear staining was performed with a 1:500 DAPI solution for 5 minutes at room temperature. Slides were then washed in TBS for 10 minutes, dipped in ddH₂O for 10 seconds, and air-dried with pressurized air. The prepared slides were subjected to fluorescence microscopy using a Zeiss Axioscan Z1 slide scanner. Autofocus was performed on the DAPI channel, and whole-slide imaging was acquired at fluorescence channels 488 nm, 555 nm, and 647 nm.

### Data preprocessing using gutenTAG

After acquisition, MALDI-IHC data were converted to .i*mzML* and *ibd* formats, then imported into the gutenTAG pipeline either manually or via the B-Fabric^42^ platform from the Functional Genomics Center Zürich. R version 4.4.1 was used for gutenTAG processing in B-fabric. R version 4.3.1 is used for analysis.

#### Preprocessing

Data normalization was performed using the Total Ion Current (TIC) method to adjust for variations across pixels while preserving biological variability. Spectral smoothing was applied using a Gaussian filter to enhance signal quality by reducing noise, and baseline reduction was achieved through a local minimum approach, where the baseline was estimated within small m/z windows and subtracted to improve peak detection. Default parameters from the Cardinal R package were used for all preprocessing steps.

#### Metapeak Generation

Metapeak construction was carried out by summing the number of detected peaks to generate a counts histogram, followed by Gaussian smoothing (sd=1) to overlap isotopic peak counts. Count-based processing of MSI spectra has previously been described for establishing consensus peak location^31^. A watershed algorithm from the R package EBImage was then applied to the smoothed counts to cluster isotopic peaks together and segment isotopic envelopes. Any peak present in fewer than 1% of pixels was filtered out, ensuring only biologically relevant peaks were retained. Peaks that are segmented together are termed ‘Metapeaks’. The Metapeak center is the m/z value with the highest peak counts within the limits of that Metapeak; the center acts as a consensus value for the location of the majority of the peaks. Metapeak centers are determined automatically, while metapeak limits can be determined automatically by gutenTAG or set manually by the user. We found that constraining metapeak limits to 0.5m/z either side of the metapeak center helps to mitigate batch effects in multi-sample analyses. Finally, Metapeaks are assigned to the reference panel of mass tags; Metapeak centers within 1m/z of a mass tag in the panel are annotated with that tag label, while Metapeaks that are outside of this association threshold are stored as ‘untargeted Metapeaks’.

#### Mean-Variance Calculation

The mean-variance relationship was examined by regressing marker variance over the mean intensity. The residuals of this regression are termed the “corrected standard deviation”. This allows for accurate comparison of marker variability independent of mean intensity differences.

#### Signal-to-Noise Ratio (SNR) Calculation

A signal-noise **ratio** was derived by employing a Gaussian Mixture Model (GMM) to classify pixel intensities as either signal or noise. SNR is defined here as the ratio between the mean intensity of the signal and noise clusters. The R package mclust was used for GMM calculation. Otsu’s method is used for determining the threshold for total ion current (TIC) histogram.

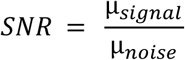

#### Geary’s C Calculation

Geary’s C was computed to quantify spatial autocorrelation, calculated as:

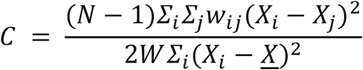

Where *w*_*ij*_ is a spatial weight matrix, *X*_*i*_ is the intensity value in pixel i, *X*_*j*_ is the intensity value in pixel j, <u>*X*</u> is the mean intensity value and W is the sum of all values in the spatial weight matrix.

#### Trashcan plot

Summarizing the data from SNR score and Geary’s C score, a trashcan plot was generated by plotting S/N separation scores against Geary’s C scores, flagging markers that failed both criteria for exclusion. Markers positioned in the lower right quadrant were identified for removal, while intermediate markers required further review. For all datasets processed in this work, mass reference lists were generated after excluding low-quality markers flagged by the gutenTAG trashcan plot. Additionally, some borderline markers were also removed to ensure high-quality downstream region segmentation.

### Downstream data analysis

#### Clustering

For single MALDI-IHC datasets, gutenTAG processed data was clustered using a k-means algorithm using a correlation distance metric. For clustering of multiple samples, each dataset was first individually log-transformed and z-scaled to stabilize variance across samples. Datasets were then concatenated and clustered together as above. In all cases, the choice of *k* was determined by computing the stability of the clustering using adjusted rand index (ARI) of the clustering across a range of *k* values. K-means clustering was run *n* times for each *k* value and a different set seed was chosen for each run to ensure random initialization of each clustering. In each case, the number of clusters was determined by choosing the *k* value with the highest mean ARI value across all runs. For the endometrial cohort, k=16 had the highest ARI value, however necrotic and vessel-containing pixels were not differentiated at this granularity, so k=19 was chosen as it had the next highest value.

#### Moran’s I analysis

To assess spatial autocorrelation of segmentation clusters within the MALDI-IHC cohort dataset, Moran’s I statistic was calculated using the spdep package in R. Spatial coordinate data for each segmentation region from each sample were used in the calculation. The resulting Moran’s I values were plotted using ggplot2 to visualize spatial autocorrelation levels across cohort samples and compiled into a summary table recording the Moran’s I value for each segmentation region in each sample.

## Data availability

All raw data and all antibody validation data will be available upon publication.

## Code availability

All code will be available upon publication (https://github.com/BodenmillerGroup)

## Acknowledgements

B.B. was supported by two Swiss National Science Foundation (SNSF) grants (310030_205007, 316030_213512), a US National Institutes of Health grant (5R01DK131059-04) and the European Research Council (ERC) under the European Union’s Horizon 2020 Program under ERC grant agreement no. 866074 (“Precision Motifs”). This work was also supported by SBIR grants to AmberGen Inc. from the National Institutes of Health including the following: R44CA236097, R44AG078097 and R44MH132196. We thank the Functional Genomics Center Zurich (FGCZ), in particular Leonardo Schwarz, Dr. Christian Panse and Dr. Witold Wolski, for their contributions and for enabling gutenTAG processing to run on B-fabric, the data management framework of the FGCZ.

## Supplementary Information

### Supplementary Tables

**Supplementary Table 1.**
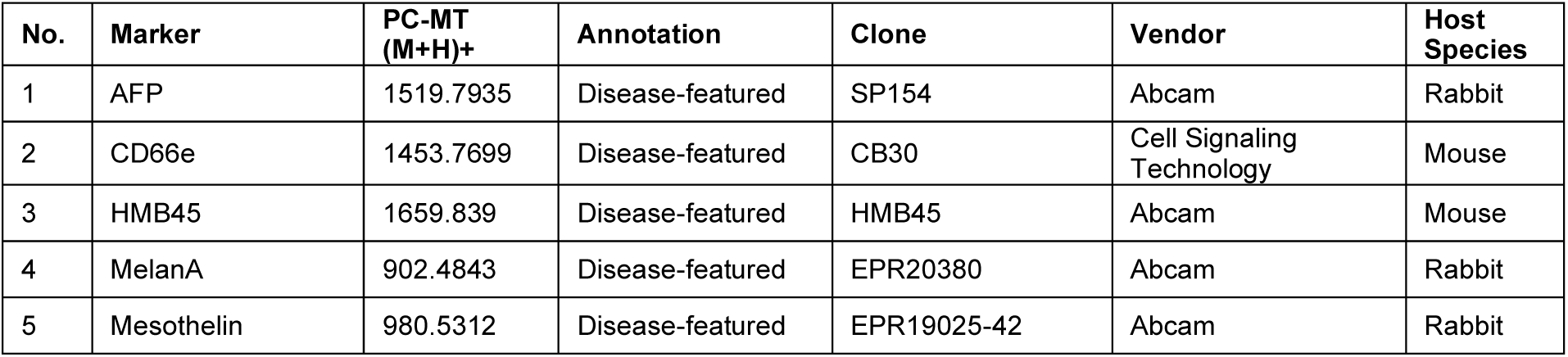
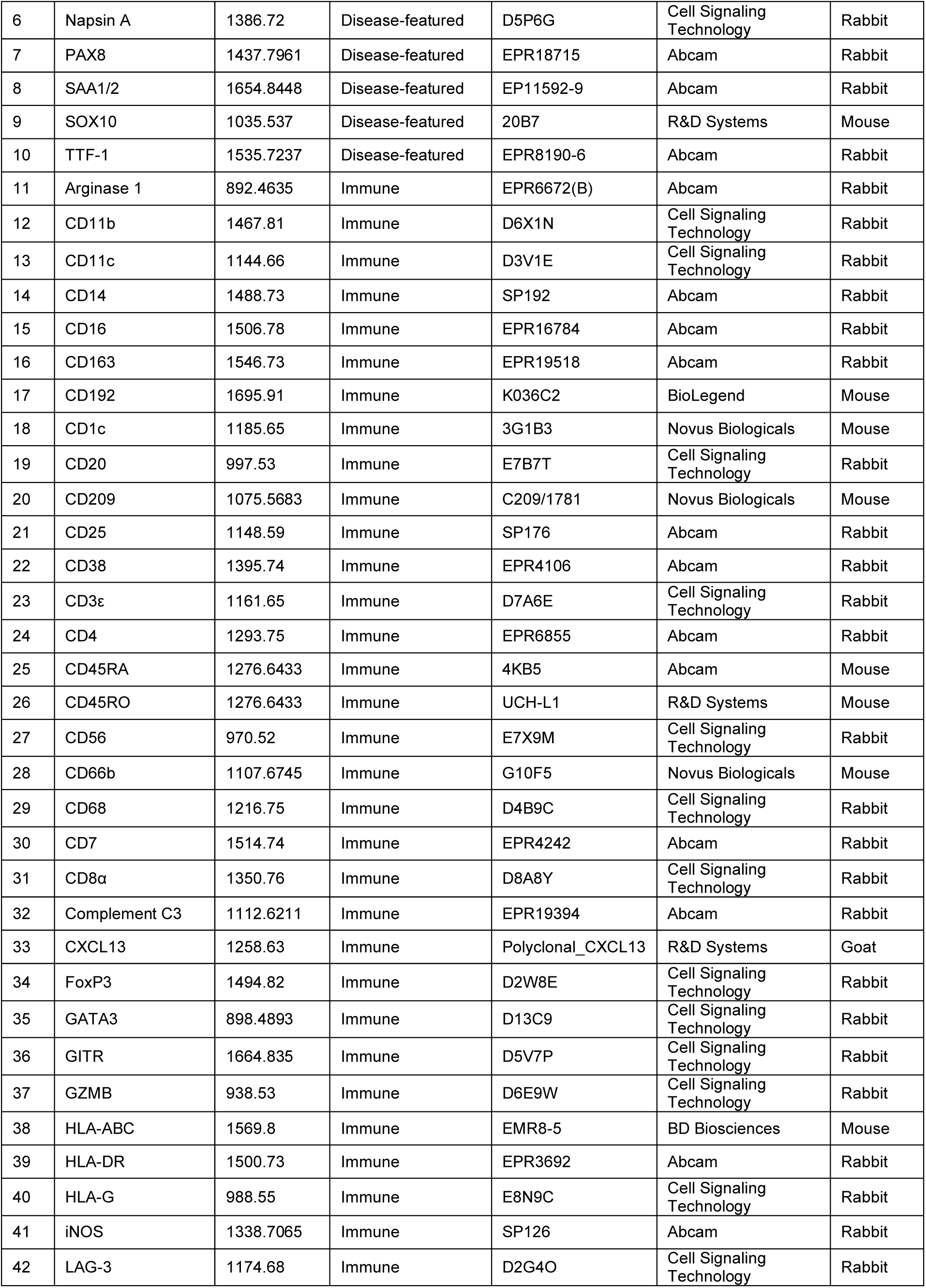
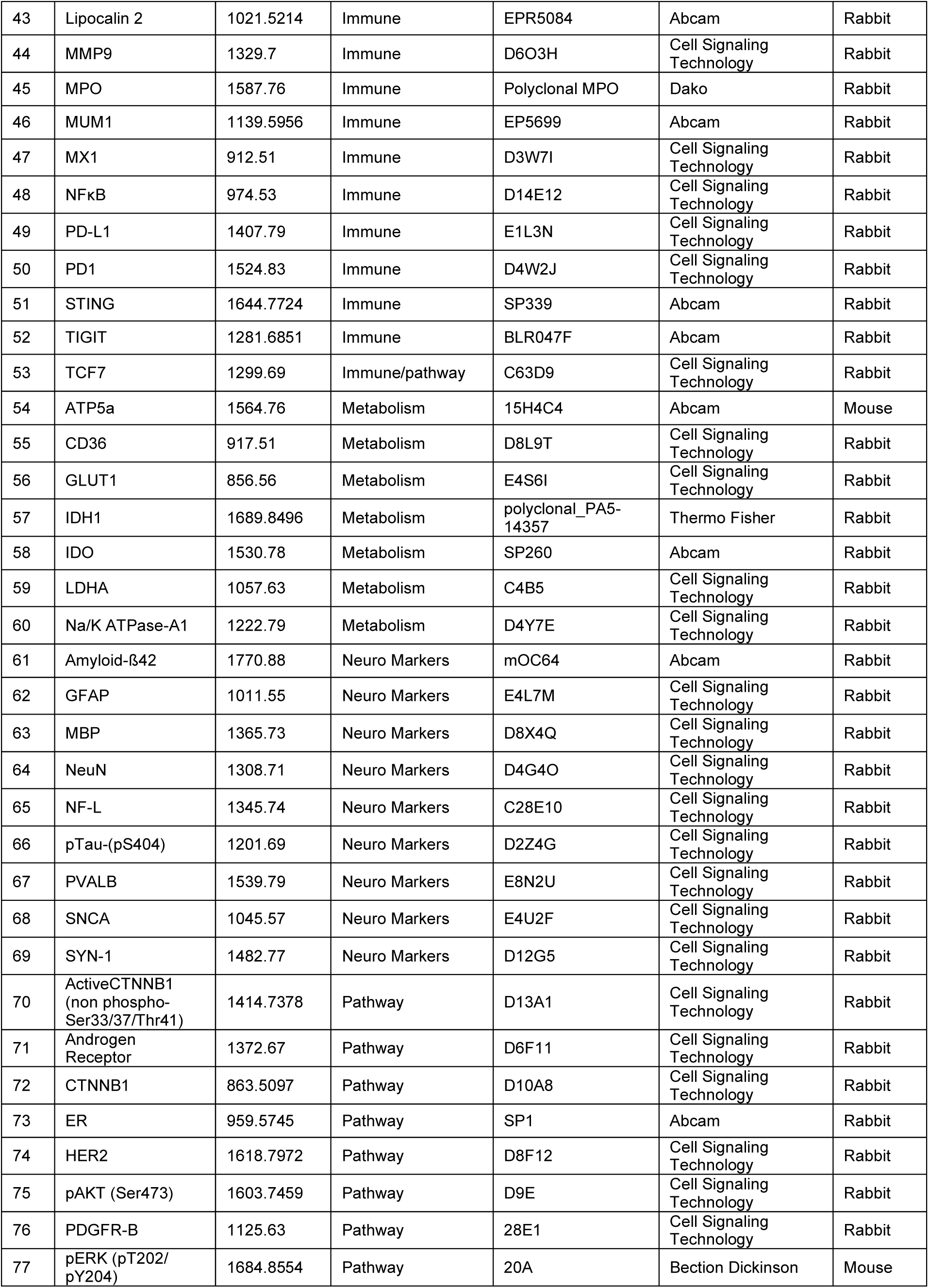
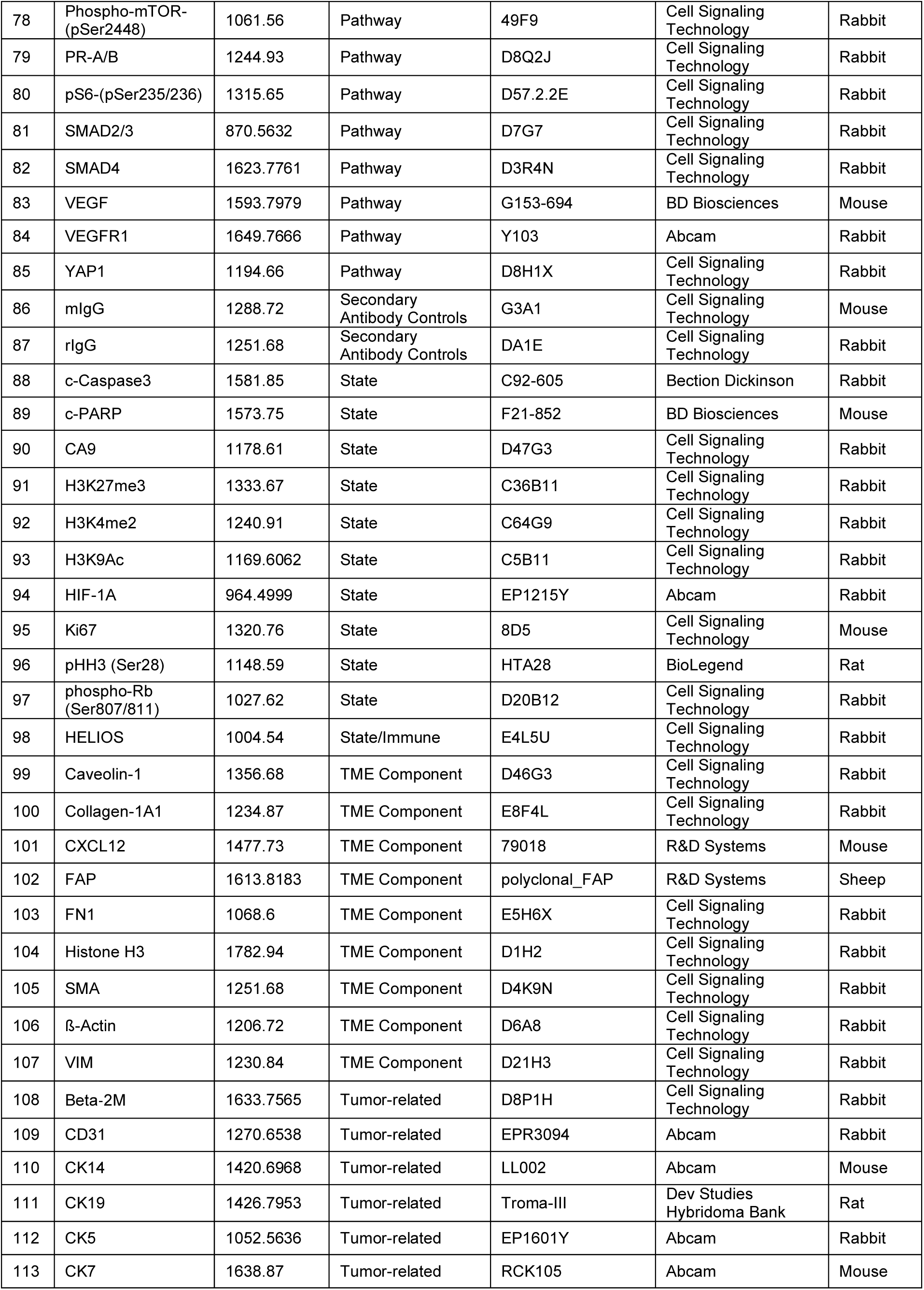
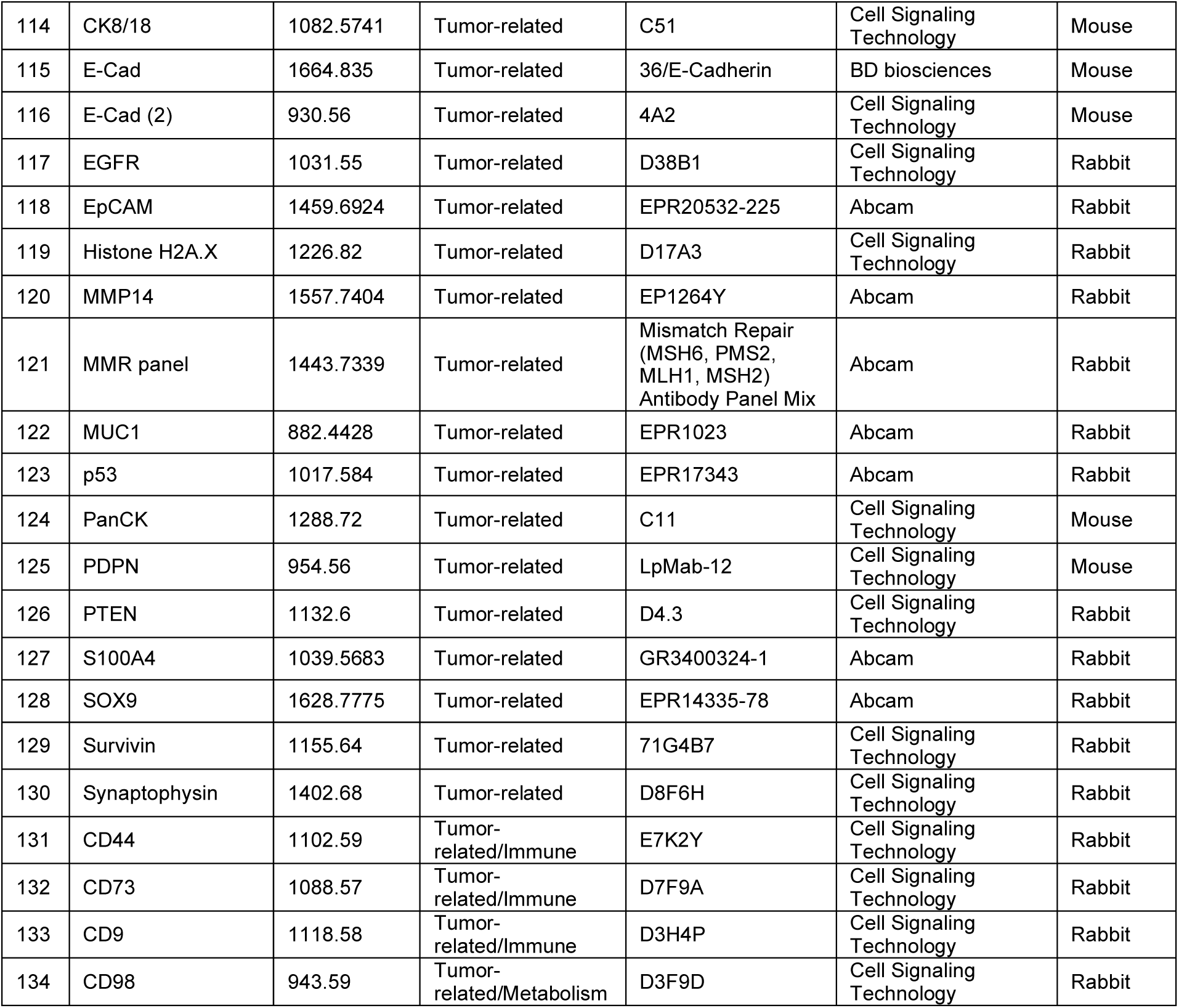
List of all 134 antibodies with clone and vendor details; two E-Cadherin clones were used, resulting in a 133-plex.

| No. | Marker | PC-MT (M+H)+ | Annotation | Clone | Vendor | Host Species |
| --- | --- | --- | --- | --- | --- | --- |
| 1 | AFP | 1519.7935 | Disease-featured | SP154 | Abcam | Rabbit |
| 2 | CD66e | 1453.7699 | Disease-featured | CB30 | Cell Signaling Technology | Mouse |
| 3 | HMB45 | 1659.839 | Disease-featured | HMB45 | Abcam | Mouse |
| 4 | MelanA | 902.4843 | Disease-featured | EPR20380 | Abcam | Rabbit |
| 5 | Mesothelin | 980.5312 | Disease-featured | EPR19025-42 | Abcam | Rabbit |
| 6 | Napsin A | 1386.72 | Disease-featured | D5P6G | Cell Signaling Technology | Rabbit |
| 7 | PAX8 | 1437.7961 | Disease-featured | EPR18715 | Abcam | Rabbit |
| 8 | SAA1/2 | 1654.8448 | Disease-featured | EP11592-9 | Abcam | Rabbit |
| 9 | SOX10 | 1035.537 | Disease-featured | 20B7 | R&D Systems | Mouse |
| 10 | TTF-1 | 1535.7237 | Disease-featured | EPR8190-6 | Abcam | Rabbit |
| 11 | Arginase 1 | 892.4635 | Immune | EPR6672(B) | Abcam | Rabbit |
| 12 | CD11b | 1467.81 | Immune | D6X1N | Cell Signaling Technology | Rabbit |
| 13 | CD11c | 1144.66 | Immune | D3V1E | Cell Signaling Technology | Rabbit |
| 14 | CD14 | 1488.73 | Immune | SP192 | Abcam | Rabbit |
| 15 | CD16 | 1506.78 | Immune | EPR16784 | Abcam | Rabbit |
| 16 | CD163 | 1546.73 | Immune | EPR19518 | Abcam | Rabbit |
| 17 | CD192 | 1695.91 | Immune | K036C2 | BioLegend | Mouse |
| 18 | CD1c | 1185.65 | Immune | 3G1B3 | Novus Biologicals | Mouse |
| 19 | CD20 | 997.53 | Immune | E7B7T | Cell Signaling Technology | Rabbit |
| 20 | CD209 | 1075.5683 | Immune | C209/1781 | Novus Biologicals | Mouse |
| 21 | CD25 | 1148.59 | Immune | SP176 | Abcam | Rabbit |
| 22 | CD38 | 1395.74 | Immune | EPR4106 | Abcam | Rabbit |
| 23 | CD3ε | 1161.65 | Immune | D7A6E | Cell Signaling Technology | Rabbit |
| 24 | CD4 | 1293.75 | Immune | EPR6855 | Abcam | Rabbit |
| 25 | CD45RA | 1276.6433 | Immune | 4KB5 | Abcam | Mouse |
| 26 | CD45RO | 1276.6433 | Immune | UCH-L1 | R&D Systems | Mouse |
| 27 | CD56 | 970.52 | Immune | E7X9M | Cell Signaling Technology | Rabbit |
| 28 | CD66b | 1107.6745 | Immune | G10F5 | Novus Biologicals | Mouse |
| 29 | CD68 | 1216.75 | Immune | D4B9C | Cell Signaling Technology | Rabbit |
| 30 | CD7 | 1514.74 | Immune | EPR4242 | Abcam | Rabbit |
| 31 | CD8α | 1350.76 | Immune | D8A8Y | Cell Signaling Technology | Rabbit |
| 32 | Complement C3 | 1112.6211 | Immune | EPR19394 | Abcam | Rabbit |
| 33 | CXCL13 | 1258.63 | Immune | Polyclonal_CXCL13 | R&D Systems | Goat |
| 34 | FoxP3 | 1494.82 | Immune | D2W8E | Cell Signaling Technology | Rabbit |
| 35 | GATA3 | 898.4893 | Immune | D13C9 | Cell Signaling Technology | Rabbit |
| 36 | GITR | 1664.835 | Immune | D5V7P | Cell Signaling Technology | Rabbit |
| 37 | GZMB | 938.53 | Immune | D6E9W | Cell Signaling Technology | Rabbit |
| 38 | HLA-ABC | 1569.8 | Immune | EMR8-5 | BD Biosciences | Mouse |
| 39 | HLA-DR | 1500.73 | Immune | EPR3692 | Abcam | Rabbit |
| 40 | HLA-G | 988.55 | Immune | E8N9C | Cell Signaling Technology | Rabbit |
| 41 | iNOS | 1338.7065 | Immune | SP126 | Abcam | Rabbit |
| 42 | LAG-3 | 1174.68 | Immune | D2G4O | Cell Signaling Technology | Rabbit |
| 43 | Lipocalin 2 | 1021.5214 | Immune | EPR5084 | Abcam | Rabbit |
| 44 | MMP9 | 1329.7 | Immune | D6O3H | Cell Signaling Technology | Rabbit |
| 45 | MPO | 1587.76 | Immune | Polyclonal MPO | Dako | Rabbit |
| 46 | MUM1 | 1139.5956 | Immune | EP5699 | Abcam | Rabbit |
| 47 | MX1 | 912.51 | Immune | D3W7I | Cell Signaling Technology | Rabbit |
| 48 | NFκB | 974.53 | Immune | D14E12 | Cell Signaling Technology | Rabbit |
| 49 | PD-L1 | 1407.79 | Immune | E1L3N | Cell Signaling Technology | Rabbit |
| 50 | PD1 | 1524.83 | Immune | D4W2J | Cell Signaling Technology | Rabbit |
| 51 | STING | 1644.7724 | Immune | SP339 | Abcam | Rabbit |
| 52 | TIGIT | 1281.6851 | Immune | BLR047F | Abcam | Rabbit |
| 53 | TCF7 | 1299.69 | Immune/pathway | C63D9 | Cell Signaling Technology | Rabbit |
| 54 | ATP5a | 1564.76 | Metabolism | 15H4C4 | Abcam | Mouse |
| 55 | CD36 | 917.51 | Metabolism | D8L9T | Cell Signaling Technology | Rabbit |
| 56 | GLUT1 | 856.56 | Metabolism | E4S6I | Cell Signaling Technology | Rabbit |
| 57 | IDH1 | 1689.8496 | Metabolism | polyclonal_PA5-14357 | Thermo Fisher | Rabbit |
| 58 | IDO | 1530.78 | Metabolism | SP260 | Abcam | Rabbit |
| 59 | LDHA | 1057.63 | Metabolism | C4B5 | Cell Signaling Technology | Rabbit |
| 60 | Na/K ATPase-A1 | 1222.79 | Metabolism | D4Y7E | Cell Signaling Technology | Rabbit |
| 61 | Amyloid-β42 | 1770.88 | Neuro Markers | mOC64 | Abcam | Rabbit |
| 62 | GFAP | 1011.55 | Neuro Markers | E4L7M | Cell Signaling Technology | Rabbit |
| 63 | MBP | 1365.73 | Neuro Markers | D8X4Q | Cell Signaling Technology | Rabbit |
| 64 | NeuN | 1308.71 | Neuro Markers | D4G4O | Cell Signaling Technology | Rabbit |
| 65 | NF-L | 1345.74 | Neuro Markers | C28E10 | Cell Signaling Technology | Rabbit |
| 66 | pTau-(pS404) | 1201.69 | Neuro Markers | D2Z4G | Cell Signaling Technology | Rabbit |
| 67 | PVALB | 1539.79 | Neuro Markers | E8N2U | Cell Signaling Technology | Rabbit |
| 68 | SNCA | 1045.57 | Neuro Markers | E4U2F | Cell Signaling Technology | Rabbit |
| 69 | SYN-1 | 1482.77 | Neuro Markers | D12G5 | Cell Signaling Technology | Rabbit |
| 70 | ActiveCTNNB1 (non phospho-Ser33/37/Thr41) | 1414.7378 | Pathway | D13A1 | Cell Signaling Technology | Rabbit |
| 71 | Androgen Receptor | 1372.67 | Pathway | D6F11 | Cell Signaling Technology | Rabbit |
| 72 | CTNNB1 | 863.5097 | Pathway | D10A8 | Cell Signaling Technology | Rabbit |
| 73 | ER | 959.5745 | Pathway | SP1 | Abcam | Rabbit |
| 74 | HER2 | 1618.7972 | Pathway | D8F12 | Cell Signaling Technology | Rabbit |
| 75 | pAKT (Ser473) | 1603.7459 | Pathway | D9E | Cell Signaling Technology | Rabbit |
| 76 | PDGFR-B | 1125.63 | Pathway | 28E1 | Cell Signaling Technology | Rabbit |
| 77 | pERK (pT202/pY204) | 1684.8554 | Pathway | 20A | Bection Dickinson | Mouse |
| 78 | Phospho-mTOR-(pSer2448) | 1061.56 | Pathway | 49F9 | Cell Signaling Technology | Rabbit |
| 79 | PR-A/B | 1244.93 | Pathway | D8Q2J | Cell Signaling Technology | Rabbit |
| 80 | pS6-(pSer235/236) | 1315.65 | Pathway | D57.2.2E | Cell Signaling Technology | Rabbit |
| 81 | SMAD2/3 | 870.5632 | Pathway | D7G7 | Cell Signaling Technology | Rabbit |
| 82 | SMAD4 | 1623.7761 | Pathway | D3R4N | Cell Signaling Technology | Rabbit |
| 83 | VEGF | 1593.7979 | Pathway | G153-694 | BD Biosciences | Mouse |
| 84 | VEGFR1 | 1649.7666 | Pathway | Y103 | Abcam | Rabbit |
| 85 | YAP1 | 1194.66 | Pathway | D8H1X | Cell Signaling Technology | Rabbit |
| 86 | mIgG | 1288.72 | Secondary Antibody Controls | G3A1 | Cell Signaling Technology | Mouse |
| 87 | rlgG | 1251.68 | Secondary Antibody Controls | DA1E | Cell Signaling Technology | Rabbit |
| 88 | c-Caspase3 | 1581.85 | State | C92-605 | Bection Dickinson | Rabbit |
| 89 | c-PARP | 1573.75 | State | F21-852 | BD Biosciences | Mouse |
| 90 | CA9 | 1178.61 | State | D47G3 | Cell Signaling Technology | Rabbit |
| 91 | H3K27me3 | 1333.67 | State | C36B11 | Cell Signaling Technology | Rabbit |
| 92 | H3K4me2 | 1240.91 | State | C64G9 | Cell Signaling Technology | Rabbit |
| 93 | H3K9Ac | 1169.6062 | State | C5B11 | Cell Signaling Technology | Rabbit |
| 94 | HIF-1A | 964.4999 | State | EP1215Y | Abcam | Rabbit |
| 95 | Ki67 | 1320.76 | State | 8D5 | Cell Signaling Technology | Mouse |
| 96 | pHH3 (Ser28) | 1148.59 | State | HTA28 | BioLegend | Rat |
| 97 | phospho-Rb (Ser807/811) | 1027.62 | State | D20B12 | Cell Signaling Technology | Rabbit |
| 98 | HELIOS | 1004.54 | State/Immune | E4L5U | Cell Signaling Technology | Rabbit |
| 99 | Caveolin-1 | 1356.68 | TME Component | D46G3 | Cell Signaling Technology | Rabbit |
| 100 | Collagen-1A1 | 1234.87 | TME Component | E8F4L | Cell Signaling Technology | Rabbit |
| 101 | CXCL12 | 1477.73 | TME Component | 79018 | R&D Systems | Mouse |
| 102 | FAP | 1613.8183 | TME Component | polyclonal_FAP | R&D Systems | Sheep |
| 103 | FN1 | 1068.6 | TME Component | E5H6X | Cell Signaling Technology | Rabbit |
| 104 | Histone H3 | 1782.94 | TME Component | D1H2 | Cell Signaling Technology | Rabbit |
| 105 | SMA | 1251.68 | TME Component | D4K9N | Cell Signaling Technology | Rabbit |
| 106 | β-Actin | 1206.72 | TME Component | D6A8 | Cell Signaling Technology | Rabbit |
| 107 | VIM | 1230.84 | TME Component | D21H3 | Cell Signaling Technology | Rabbit |
| 108 | Beta-2M | 1633.7565 | Tumor-related | D8P1H | Cell Signaling Technology | Rabbit |
| 109 | CD31 | 1270.6538 | Tumor-related | EPR3094 | Abcam | Rabbit |
| 110 | CK14 | 1420.6968 | Tumor-related | LL002 | Abcam | Mouse |
| 111 | CK19 | 1426.7953 | Tumor-related | Troma-III | Dev Studies Hybridoma Bank | Rat |
| 112 | CK5 | 1052.5636 | Tumor-related | EP1601Y | Abcam | Rabbit |
| 113 | CK7 | 1638.87 | Tumor-related | RCK105 | Abcam | Mouse |
| 114 | CK8/18 | 1082.5741 | Tumor-related | C51 | Cell Signaling Technology | Mouse |
| 115 | E-Cad | 1664.835 | Tumor-related | 36/E-Cadherin | BD biosciences | Mouse |
| 116 | E-Cad (2) | 930.56 | Tumor-related | 4A2 | Cell Signaling Technology | Mouse |
| 117 | EGFR | 1031.55 | Tumor-related | D38B1 | Cell Signaling Technology | Rabbit |
| 118 | EpCAM | 1459.6924 | Tumor-related | EPR20532-225 | Abcam | Rabbit |
| 119 | Histone H2A.X | 1226.82 | Tumor-related | D17A3 | Cell Signaling Technology | Rabbit |
| 120 | MMP14 | 1557.7404 | Tumor-related | EP1264Y | Abcam | Rabbit |
| 121 | MMR panel | 1443.7339 | Tumor-related | Mismatch Repair (MSH6, PMS2, MLH1, MSH2) Antibody Panel Mix | Abcam | Rabbit |
| 122 | MUC1 | 882.4428 | Tumor-related | EPR1023 | Abcam | Rabbit |
| 123 | p53 | 1017.584 | Tumor-related | EPR17343 | Abcam | Rabbit |
| 124 | PanCK | 1288.72 | Tumor-related | C11 | Cell Signaling Technology | Mouse |
| 125 | PDPN | 954.56 | Tumor-related | LpMab-12 | Cell Signaling Technology | Mouse |
| 126 | PTEN | 1132.6 | Tumor-related | D4.3 | Cell Signaling Technology | Rabbit |
| 127 | S100A4 | 1039.5683 | Tumor-related | GR3400324-1 | Abcam | Rabbit |
| 128 | SOX9 | 1628.7775 | Tumor-related | EPR14335-78 | Abcam | Rabbit |
| 129 | Survivin | 1155.64 | Tumor-related | 71G4B7 | Cell Signaling Technology | Rabbit |
| 130 | Synaptophysin | 1402.68 | Tumor-related | D8F6H | Cell Signaling Technology | Rabbit |
| 131 | CD44 | 1102.59 | Tumor-related/Immune | E7K2Y | Cell Signaling Technology | Rabbit |
| 132 | CD73 | 1088.57 | Tumor-related/Immune | D7F9A | Cell Signaling Technology | Rabbit |
| 133 | CD9 | 1118.58 | Tumor-related/Immune | D3H4P | Cell Signaling Technology | Rabbit |
| 134 | CD98 | 943.59 | Tumor-related/Metabolism | D3F9D | Cell Signaling Technology | Rabbit |

**Supplementary Table 2.** The 120-plex panel for high-plex experiments, including staining concentrations for each marker.

| No. | Name | PC-MT (M+H)+ | Annotation | Staining Concentration (µg/mL) |
| --- | --- | --- | --- | --- |
| 1 | MelanA | 902.4843 | Disease-featured | 0.5 |
| 2 | Mesothelin | 980.5312 | Disease-featured | 1 |
| 3 | SOX10 | 1035.537 | Disease-featured | 2 |
| 4 | Napsin A | 1386.72 | Disease-featured | 1 |
| 5 | PAX8 | 1437.7961 | Disease-featured | 1 |
| 6 | CD66e | 1453.7699 | Disease-featured | 1 |
| 7 | AFP | 1519.7935 | Disease-featured | 1 |
| 8 | TTF-1 | 1535.7237 | Disease-featured | 1 |
| 9 | SAA1/2 | 1654.8448 | Disease-featured | 1 |
| 10 | HMB45 | 1659.839 | Disease-featured | 1 |
| 11 | Arginase 1 | 892.4635 | Immune | 0.5 |
| 12 | GATA3 | 898.4893 | Immune | 0.5 |
| 13 | MX1 | 912.51 | Immune | 1 |
| 14 | GZMB | 938.53 | Immune | 1 |
| 15 | CD56 | 970.52 | Immune | 1 |
| 16 | NFκB | 974.53 | Immune | 1 |
| 17 | HLA-G | 988.55 | Immune | 1 |
| 18 | CD20 | 997.53 | Immune | 0.5 |
| 19 | Lipocalin 2 | 1021.5214 | Immune | 1 |
| 20 | CD209 | 1075.5683 | Immune | 0.5 |
| 21 | CD66b | 1107.6745 | Immune | 1 |
| 22 | Complement C3 | 1112.6211 | Immune | 1 |
| 23 | MUM1 | 1139.5956 | Immune | 2 |
| 24 | CD11c | 1144.66 | Immune | 1 |
| 25 | CD25 | 1148.59 | Immune | 2 |
| 26 | CD3ε | 1161.65 | Immune | 1 |
| 27 | LAG-3 | 1174.68 | Immune | 2 |
| 28 | CD1c | 1185.65 | Immune | 1 |
| 29 | CD68 | 1216.75 | Immune | 1 |
| 30 | CXCL13 | 1258.63 | Immune | 1 |
| 31 | CD45RA | 1276.6433 | Immune | 1 |
| 32 | CD45RO | 1276.6433 | Immune | 1 |
| 33 | TIGIT | 1281.6851 | Immune | 1 |
| 34 | CD4 | 1293.75 | Immune | 1 |
| 35 | MMP9 | 1329.7 | Immune | 1 |
| 36 | iNOS | 1338.7065 | Immune | 1 |
| 37 | CD8α | 1350.76 | Immune | 2 |
| 38 | CD38 | 1395.74 | Immune | 0.5 |
| 39 | PD-L1 | 1407.79 | Immune | 1 |
| 40 | CD11b | 1467.81 | Immune | 0.5 |
| 41 | CD14 | 1488.73 | Immune | 1 |
| 42 | FoxP3 | 1494.82 | Immune | 1 |
| 43 | HLA-DR | 1500.73 | Immune | 0.5 |
| 44 | CD16 | 1506.78 | Immune | 1 |
| 45 | CD7 | 1514.74 | Immune | 0.5 |
| 46 | PD1 | 1524.83 | Immune | 1 |
| 47 | CD163 | 1546.73 | Immune | 1 |
| 48 | HLA-ABC | 1569.8 | Immune | 0.5 |
| 49 | MPO | 1587.76 | Immune | 1 |
| 50 | STING | 1644.7724 | Immune | 1 |
| 51 | CD192 | 1695.91 | Immune | 1 |
| 52 | TCF7 | 1299.69 | Immune/pathway | 1 |
| 53 | GLUT1 | 856.56 | Metabolism | 1 |
| 54 | CD36 | 917.51 | Metabolism | 1 |
| 55 | LDHA | 1057.63 | Metabolism | 2 |
| 56 | Na/KATPase-A1 | 1222.79 | Metabolism | 0.5 |
| 57 | IDO | 1530.78 | Metabolism | 1 |
| 58 | ATP5a | 1564.76 | Metabolism | 1 |
| 59 | IDH1 | 1689.8496 | Metabolism | 1 |
| 60 | CTNNB1 | 863.5097 | Pathway | 0.5 |
| 61 | SMAD2/3 | 870.5632 | Pathway | 1 |
| 62 | ER | 959.5745 | Pathway | 1 |
| 63 | Phospho-mTOR-(pSer2448) | 1061.56 | Pathway | 1 |
| 64 | PDGFRB | 1125.63 | Pathway | 1 |
| 65 | YAP1 | 1194.66 | Pathway | 1 |
| 66 | PR-A/B | 1244.93 | Pathway | 1 |
| 67 | pS6-(pSer235/236) | 1315.65 | Pathway | 1 |
| 68 | Androgen Receptor | 1372.67 | Pathway | 1 |
| 69 | ActiveCTNNB1 | 1414.7378 | Pathway | 1 |
| 70 | VEGF | 1593.7979 | Pathway | 1 |
| 71 | pAKT (Ser473) | 1603.7459 | Pathway | 1 |
| 72 | HER2 | 1618.7972 | Pathway | 1 |
| 73 | SMAD4 | 1623.7761 | Pathway | 1 |
| 74 | VEGFR1 | 1649.7666 | Pathway | 1 |
| 75 | pERK | 1684.8554 | Pathway | 1 |
| 76 | HIF-1A | 964.4999 | State | 0.5 |
| 77 | phospho-Rb (Ser807/811) | 1027.62 | State | 1 |
| 78 | H3K9Ac | 1169.6062 | State | 1 |
| 79 | CA9 | 1178.61 | State | 1 |
| 80 | H3K4me2 | 1240.91 | State | 1 |
| 81 | Ki67 | 1320.76 | State | 1 |
| 82 | H3K27me3 | 1333.67 | State | 0.5 |
| 83 | c-PARP | 1573.75 | State | 1 |
| 84 | c-Caspase3 | 1581.85 | State | 1 |
| 85 | HELIOS | 1004.54 | State/Immune | 2 |
| 86 | FN1 | 1068.6 | TME Component | 1 |
| 87 | Beta-Actin | 1206.72 | TME Component | 0.5 |
| 88 | VIM | 1230.84 | TME Component | 0.5 |
| 89 | Collagen-1A1 | 1234.87 | TME Component | 1 |
| 90 | SMA | 1251.68 | TME Component | 0.5 |
| 91 | Caveolin-1 | 1356.68 | TME Component | 0.5 |
| 92 | CXCL12 | 1477.73 | TME Component | 1 |
| 93 | FAP | 1613.8183 | TME Component | 1 |
| 94 | Histone H3 | 1782.94 | TME Component | 0.5 |
| 95 | MUC1 | 882.4428 | Tumor-related | 1 |
| 96 | PDPN | 954.56 | Tumor-related | 1 |
| 97 | p53 | 1017.584 | Tumor-related | 1 |
| 98 | EGFR | 1031.55 | Tumor-related | 1 |
| 99 | S100A4 | 1039.5683 | Tumor-related | 0.5 |
| 100 | CK5 | 1052.5636 | Tumor-related | 1 |
| 101 | CK8/18 | 1082.5741 | Tumor-related | 0.5 |
| 102 | PTEN | 1132.6 | Tumor-related | 1 |
| 103 | Survivin | 1155.64 | Tumor-related | 0.5 |
| 104 | Histone H2A.X | 1226.82 | Tumor-related | 1 |
| 105 | CD31 | 1270.6538 | Tumor-related | 1 |
| 106 | PanCK | 1288.72 | Tumor-related | 0.5 |
| 107 | Synaptophysin | 1402.68 | Tumor-related | 1 |
| 108 | CK14 | 1420.6968 | Tumor-related | 1 |
| 109 | CK19 | 1426.7953 | Tumor-related | 1 |
| 110 | MMR panel | 1443.7339 | Tumor-related | 8 |
| 111 | EpCAM | 1459.6924 | Tumor-related | 1 |
| 112 | MMP14 | 1557.7404 | Tumor-related | 1 |
| 113 | SOX9 | 1628.7775 | Tumor-related | 1 |
| 114 | Beta-2M | 1633.7565 | Tumor-related | 0.5 |
| 115 | CK7 | 1638.87 | Tumor-related | 1 |
| 116 | E-Cad | 1664.835 | Tumor-related | 1 |
| 117 | CD73 | 1088.57 | Tumor-related/Immune | 2 |
| 118 | CD44 | 1102.59 | Tumor-related/Immune | 0.5 |
| 119 | CD9 | 1118.58 | Tumor-related/Immune | 1 |
| 120 | CD98 | 943.59 | Tumor-related/Metabolism | 1 |

**Supplementary Table 3.** Markers for high concentration titration (up to 5µg/mL):

| Number | Name | PC-MT (M+H)+ |
| --- | --- | --- |
| 1 | AFP | 1519.7935 |
| 2 | Androgen Receptor | 1372.67 |
| 3 | ATP5a | 1564.76 |
| 4 | c-Caspase3 | 1581.85 |
| 5 | CD192 | 1695.91 |
| 6 | CD14 | 1488.73 |
| 7 | CD16 | 1506.78 |
| 8 | CD163 | 1546.73 |
| 9 | CD25 | 1148.59 |
| 10 | CD3 | 1161.65 |
| 11 | CD45RA | 1276.6433 |
| 12 | CD45RO | 1276.6433 |
| 13 | CK14 | 1420.6968 |
| 14 | CXCL12 | 1477.73 |
| 15 | GATA3 | 898.4893 |
| 16 | Histone H3 | 1782.94 |
| 17 | HLA-DR | 1500.73 |
| 18 | IDO | 1530.78 |
| 19 | IRF4 / MUM1 | 1139.5956 |
| 20 | LAG-3 | 1174.68 |
| 21 | LDHA | 1057.63 |
| 22 | MMP14 | 1557.7404 |
| 23 | pAKT (Ser473) | 1603.7459 |
| 24 | PD-L1 | 1407.79 |
| 25 | PD1 | 1524.83 |
| 26 | phospho-Rb (Ser807/811) | 1027.62 |
| 27 | SAA1/2 | 1654.8448 |
| 28 | SOX10 | 1035.537 |

**Supplementary Table 4.** 13-plex panel for the ovarian cancer showcase study.

| No. | Name | PC-MT (M+H)+ | Annotation | Function |
| --- | --- | --- | --- | --- |
| 1 | B2M | 1633.7565 | Tumor-related | MHC-I antigen, general tumor marker |
| 2 | CK19 | 1426.7953 | Tumor-related | Epithelial cell |
| 3 | CK8/18 | 1082.5741 | Tumor-related | Epithelial cell |
| 4 | MUC1 | 882.4428 | Tumor-related | Tumor |
| 5 | p53 | 1017.584 | Tumor-related | Tumor suppressor |
| 6 | S100A4 | 1039.5683 | Tumor-related | EMT, tumor invasion and metastasis |
| 7 | H3K9Ac | 1169.6062 | State | Transcriptional activation |
| 8 | ActiveCTNNB1 | 1414.7378 | Pathway | WNT signaling |
| 9 | CTNNB1 | 863.5097 | Pathway | WNT signaling |
| 10 | pAKT (Ser473) | 1603.7459 | Pathway | PI3K pathway |
| 11 | SMAD2/3 | 870.5632 | Pathway | TGF-beta pathway |
| 12 | Complement C3 | 1112.6211 | Immune | Innate immune response |
| 13 | Mesothelin | 980.5312 | Disease-featured | Ovarian cancer indicative |

**Supplementary Table 5.** 46-plex panel used in reproducibility study experiments.

| Number | Name | PC-MT (M+H)+ |
| --- | --- | --- |
| 1 | CK5 | 1052.5636 |
| 2 | PanCK | 1288.72 |
| 3 | MMR panel | 1443.7339 |
| 4 | S100A4 | 1039.5683 |
| 5 | CK8/18 | 1082.5741 |
| 6 | E-Cad | 1664.835 |
| 7 | Survivin | 1155.64 |
| 8 | B2M | 1633.7565 |
| 9 | EMA/MUC1 | 882.4428 |
| 10 | p53 | 1017.584 |
| 11 | CD44 | 1102.59 |
| 12 | CD9 | 1118.58 |
| 13 | CD98 | 943.59 |
| 14 | CD20 | 997.53 |
| 15 | GZMB | 938.53 |
| 16 | CD11c | 1144.66 |
| 17 | CD11b | 1467.81 |
| 18 | TIGIT | 1281.6851 |
| 19 | Complement C3 | 1112.6211 |
| 20 | CD66b | 1107.6745 |
| 21 | HLA-ABC | 1569.8 |
| 22 | HLA-DR | 1500.73 |
| 23 | CD14 | 1488.73 |
| 24 | Lipocalin 2 | 1021.5214 |
| 25 | MMP9 | 1329.7 |
| 26 | CD38 | 1395.74 |
| 27 | CD3 | 1161.65 |
| 28 | TCF7 | 1299.69 |
| 29 | Beta-Actin | 1206.72 |
| 30 | FAP | 1613.8183 |
| 31 | SMA | 1251.68 |
| 32 | Collagen-1A1 | 1234.87 |
| 33 | Caveolin-1 | 1356.68 |
| 34 | Histone H3 | 1782.94 |
| 35 | VIM | 1230.84 |
| 36 | Ki67 | 1320.76 |
| 37 | HIF-1A | 964.4999 |
| 38 | H3K9Ac | 1169.6062 |
| 39 | H3K4me2 | 1240.91 |
| 40 | H3K27me3 | 1333.67 |
| 41 | YAP1 | 1194.66 |
| 42 | SMAD2/3 | 870.5632 |
| 43 | CTNNB1 | 863.5097 |
| 44 | GLUT1 | 856.56 |
| 45 | LDHA | 1057.63 |
| 46 | IDH1 | 1689.8496 |

**Supplementary Table 6.** Refined panel for cohort analysis, with low-quality and biologically less relevant markers removed.

| No. | Name | PC-MT (M+H)+ |
| --- | --- | --- |
| 1 | CD11b | 1468.096 |
| 2 | HIF-1a | 964.729 |
| 3 | MMP9 | 1330.2 |
| 4 | pmTOR | 1061.773 |
| 5 | STING | 1645.352 |
| 6 | PDGFR-B | 1125.943 |
| 7 | CXCL13 | 1259.105 |
| 8 | CD11c | 1144.966 |
| 9 | NFKB | 974.811 |
| 10 | CD44 | 1102.903 |
| 11 | B2M | 1634.219 |
| 12 | VIM | 1231.052 |
| 13 | CD68 | 1216.939 |
| 14 | pS6 | 1315.999 |
| 15 | CD66b | 1107.972 |
| 16 | MX1 | 912.812 |
| 17 | CD66e | 1454.157 |
| 18 | SOX10 | 1035.825 |
| 19 | CD31 | 1271.027 |
| 20 | HLA-ABC | 1570.224 |
| 21 | FAP | 1614.144 |
| 22 | PDPN | 954.823 |
| 23 | CD209 | 1075.887 |
| 24 | Collagen-1A1 | 1235.085 |
| 25 | TIGIT | 1281.898 |
| 26 | Lipocalin 2 | 1021.891 |
| 27 | E-Cad | 1665.34 |
| 28 | CD38 | 1396.036 |
| 29 | CTNNB1 | 863.803 |
| 30 | CD20 | 997.954 |
| 31 | SMA | 1251.916 |
| 32 | PR-A/B | 1245.166 |
| 33 | H3K4me2 | 1241.134 |
| 34 | S100A4 | 1039.987 |
| 35 | YAP1 | 1194.935 |
| 36 | SMAD2/3 | 870.922 |
| 37 | H3K27me3 | 1333.97 |
| 38 | p53 | 1017.854 |
| 39 | CD98 | 943.865 |
| 40 | CD8 | 1351.938 |
| 41 | CA9 | 1178.98 |
| 42 | H3K9Ac | 1169.891 |
| 43 | MMR panel | 1444.076 |
| 44 | MUC1 | 882.73 |
| 45 | CD73 | 1088.774 |
| 46 | PTEN | 1132.869 |
| 47 | ActiveCTNNB1 | 1414.971 |
| 48 | CD9 | 1118.843 |
| 49 | PanCK | 1288.998 |
| 50 | Mesothelin | 980.859 |
| 51 | beta-Actin | 1206.945 |
| 52 | CK8/18 | 1082.988 |
| 53 | iNOS | 1339.054 |
| 54 | CK19 | 1427.069 |
| 55 | Na/K ATPase-A1 | 1223.075 |
| 56 | Arginase 1 | 892.869 |
| 57 | LDHA | 1057.916 |
| 58 | GLUT1 | 856.857 |
| 59 | Ki67 | 1321.083 |
| 60 | ER | 959.908 |
| 61 | pRb | 1027.847 |
| 62 | GZMB | 938.868 |
| 63 | Survivin | 1156.012 |
| 64 | EGFR | 1031.88 |
| 65 | CD4 | 1294.083 |
| 66 | Complement C3 | 1112.881 |
| 67 | TCF7 | 1300.132 |
| 68 | Caveolin-1 | 1356.68 |
| 69 | CD3E | 1161.973 |
| 70 | CD56 | 970.69 |

### Supplementary Figures

**Supplementary Figure 1.**
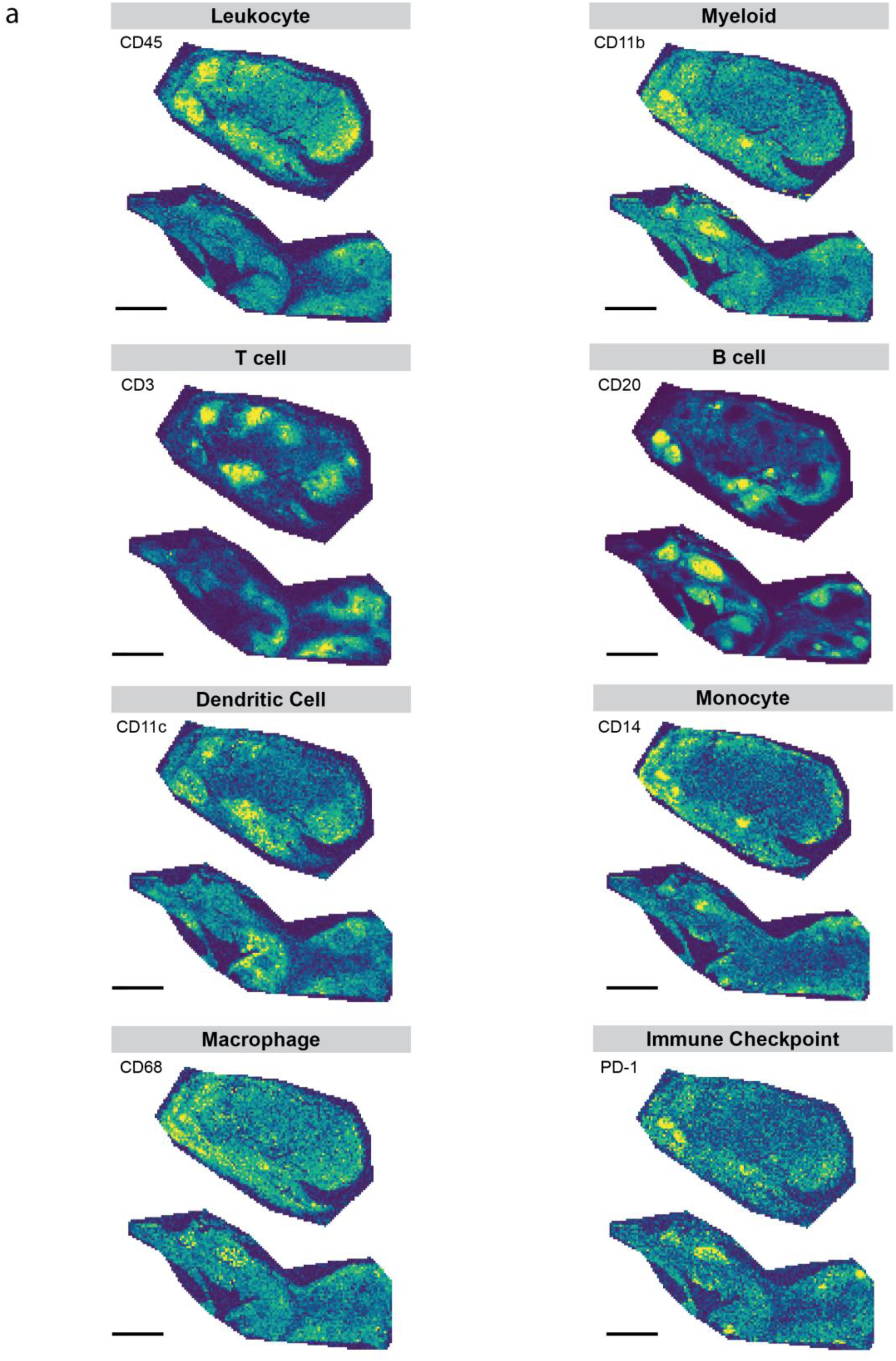
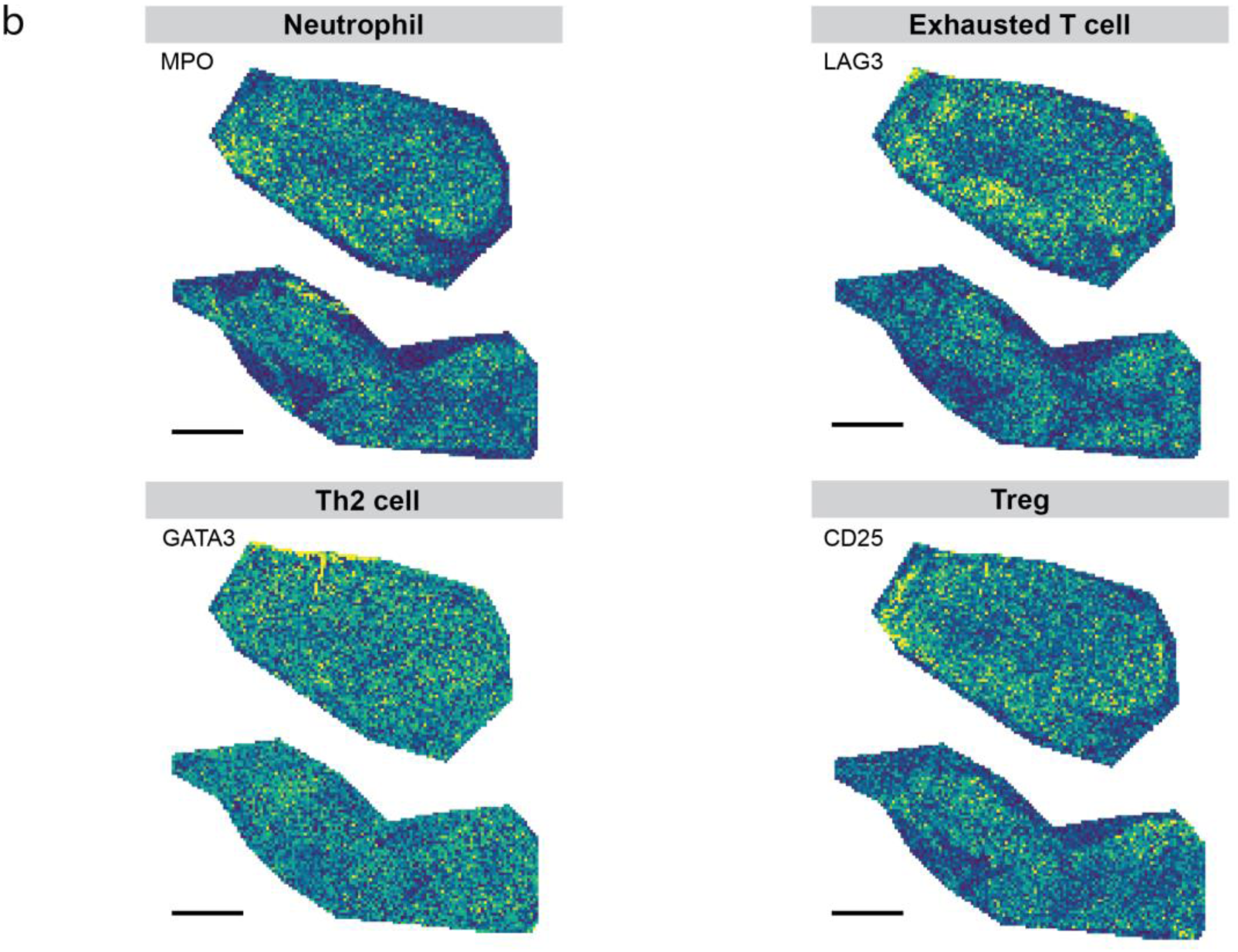
Detection of common immune markers in MALDI-IHC. **(a)** Ion images of selected immune markers following 120-plex staining of tonsil samples, acquired at 50 µm resolution. The relevant category of immune cell type is indicated in each case. Scale bar = 1 mm. **(b)** Ion images of markers that are challenging to detect.

**Supplementary Figure 2:**
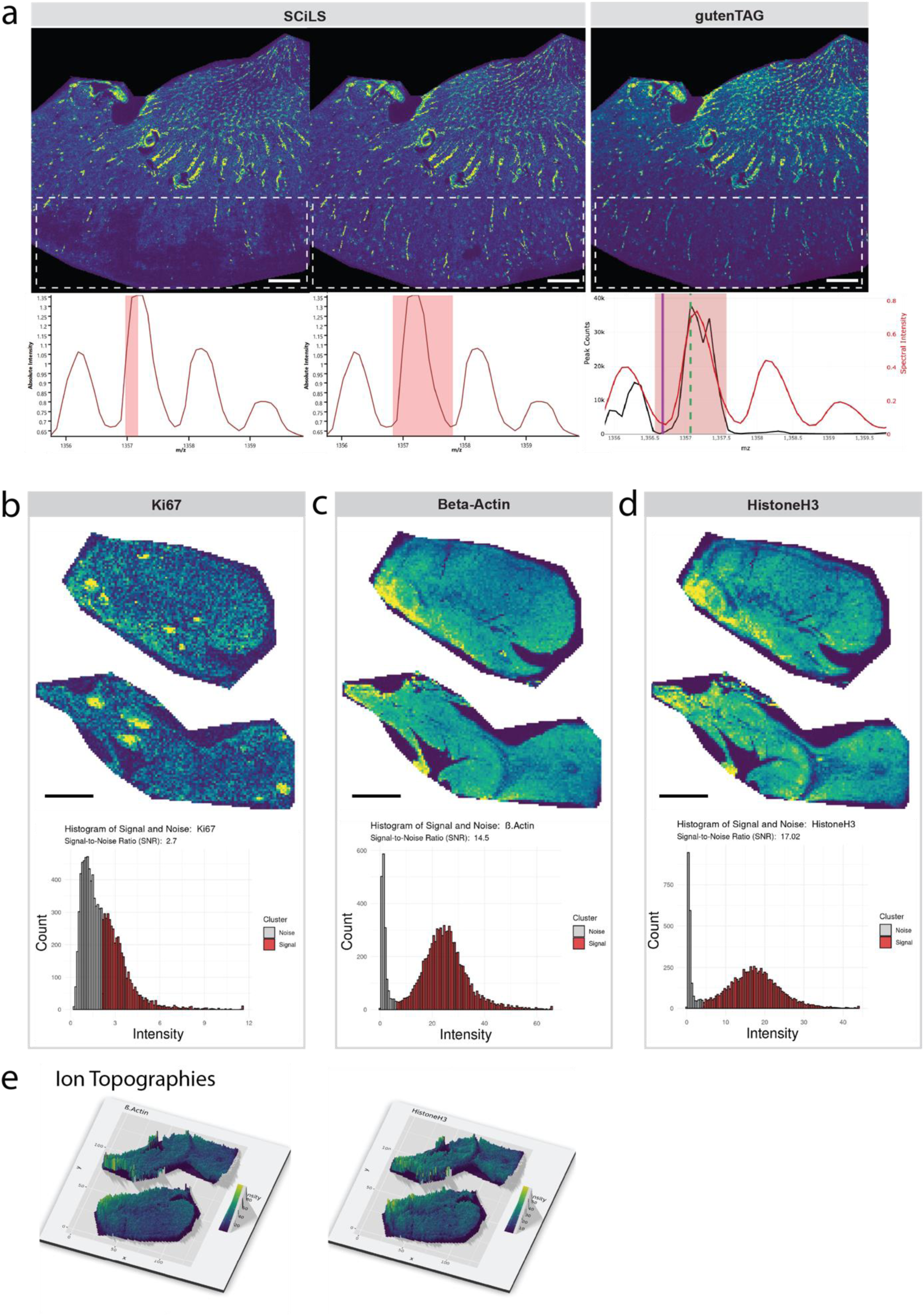
MALDI-IHC data processing with gutenTAG. **(a)** Impact of varying m/z bin widths on image fidelity, illustrated with a MALDI-IHC image of Caveolin-1 from a kidney sample at 50µm resolution. In the upper panel, the images were generated using either SCiLS Lab Pro software, with a narrow bin width (±0.100 Da, left) and a wider bin width (±0.483 Da, center), or displays the output from gutenTAG’s Metapeak processing (right image). White boxes highlight regions where the narrow bin width images show fewer signals compared to those generated with a wider bin width or Metapeak processing. The lower panel shows the corresponding zoomed-in average mass spectra; the selected bins for generating the ion images are highlighted with red boxes. The SCiLS Lab Pro traces show the mass spectra (m/z vs. intensity). In the Metapeak spectrum, the black trace is the count spectrum (m/z vs. count), while the red trace is the intensity spectrum peak (m/z vs. intensity). The green dashed line in the Metapeak spectrum marks the Metapeak peak maximum, and the purple line indicates the reference m/z. (**b-d**) Ion intensity images (upper) and intensity histograms (lower) for Ki67 (**b**), beta-Actin **(c**) and Histone H3 (**d**). In all cases, grey histograms denote noise and red histograms denote signals, as defined using a Gaussian mixture model. All scale bars = 1 mm. (**e**). Ion topographies of beta-Actin (left) and Histone H3 (right) illustrate the spatial clustering indicated by the Geary’s C scores observed in (c, d). Each topography depicts intensity gradients, with the highest intensities represented in yellow and elevated heights while areas of low intensity are depicted in dark blue and are closer to the base level.

**Supplementary Figure 3:**
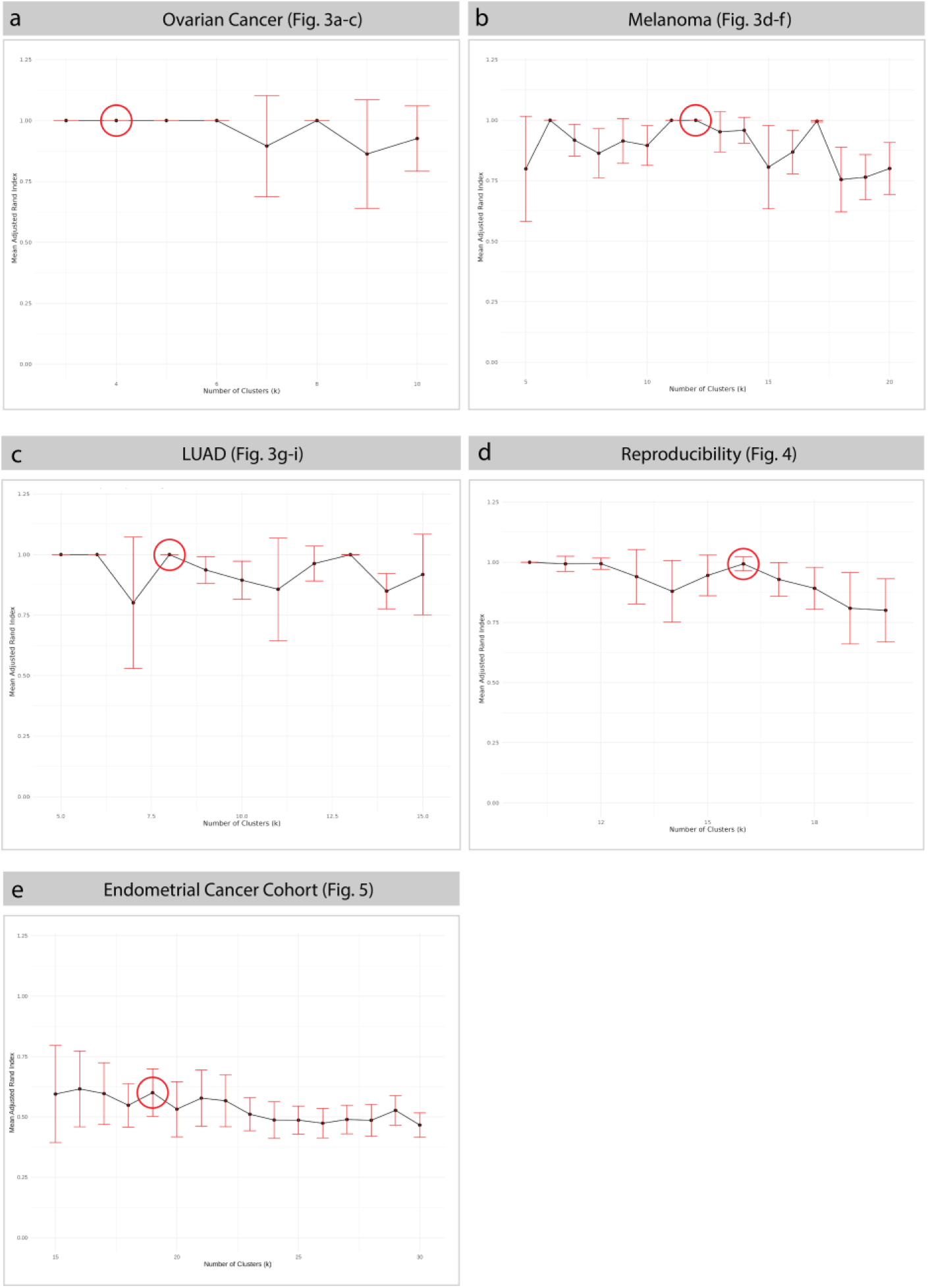
Determining the optimal number of clusters for region segmentation. **(a-e)** The plots show an adjusted rand index (ARI) stability analysis for the indicated datasets. Shown are analyses for an ovarian cancer sample (**a**), a melanoma lymph node metastasis sample (**b)**, a lung adenocarcinoma sample (**c)**, the dataset used for reproducibility analysis (**d**), and the dataset of the endometrial cancer cohort (**e)**.

**Supplementary Figure 4:**
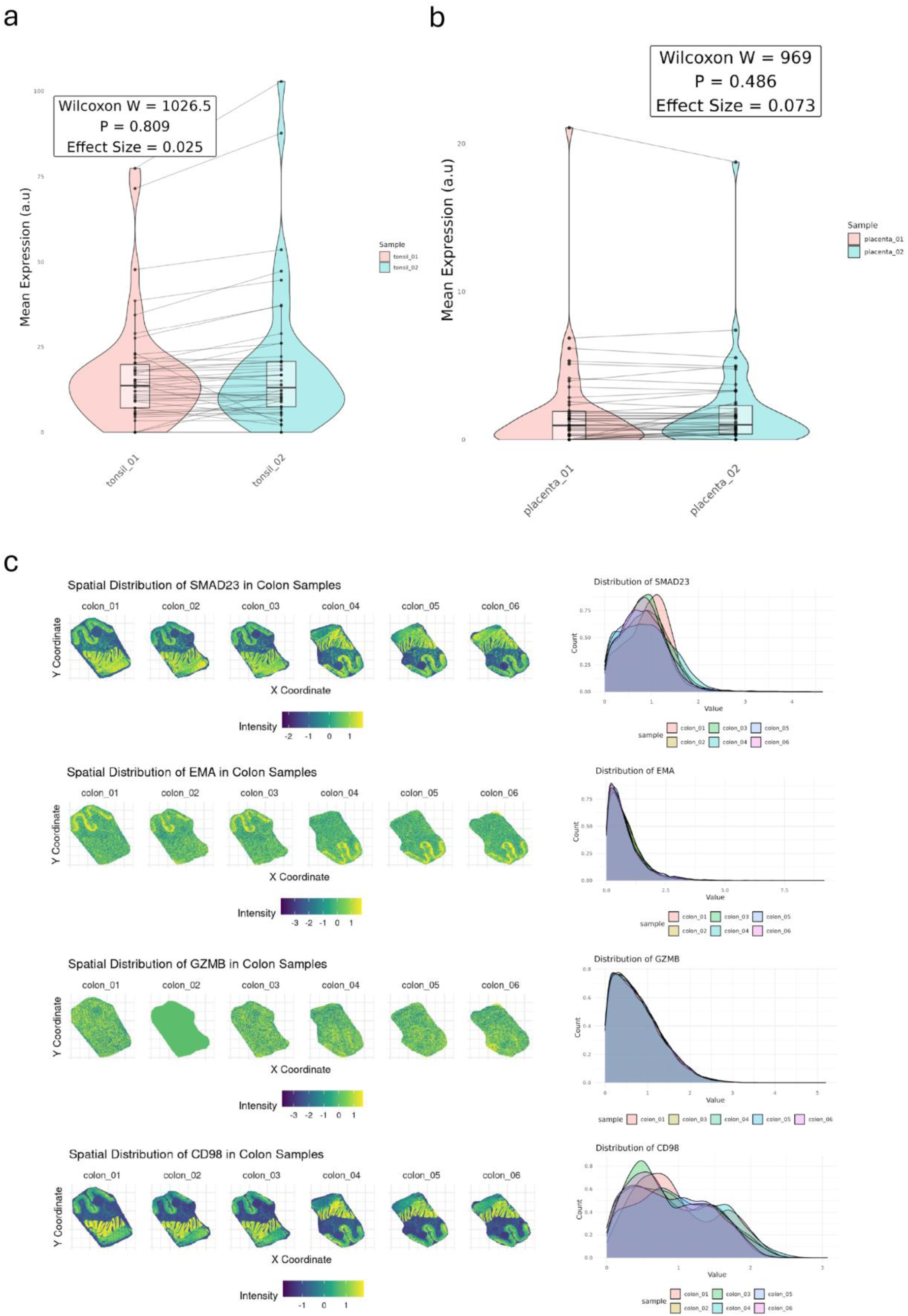
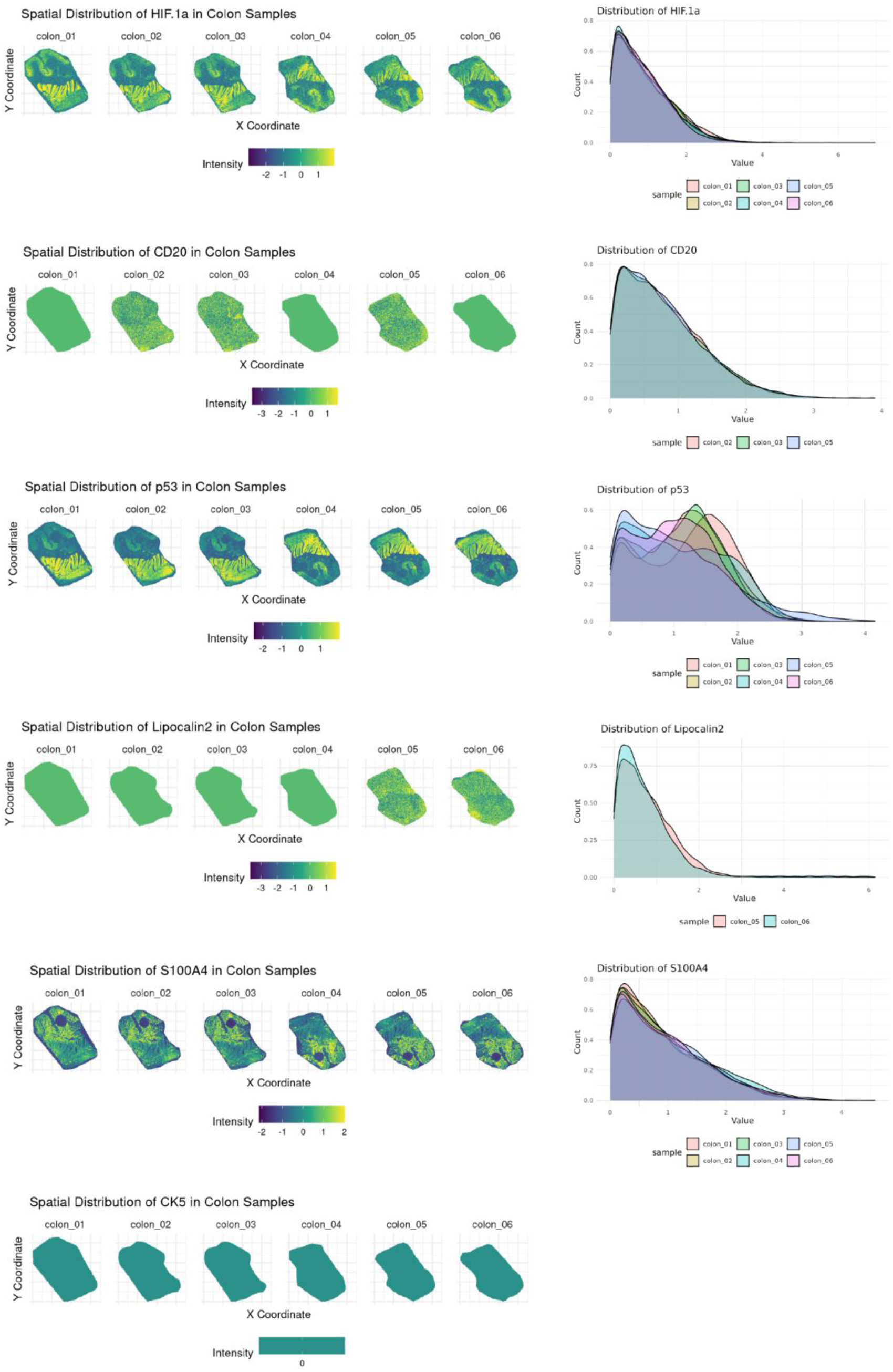
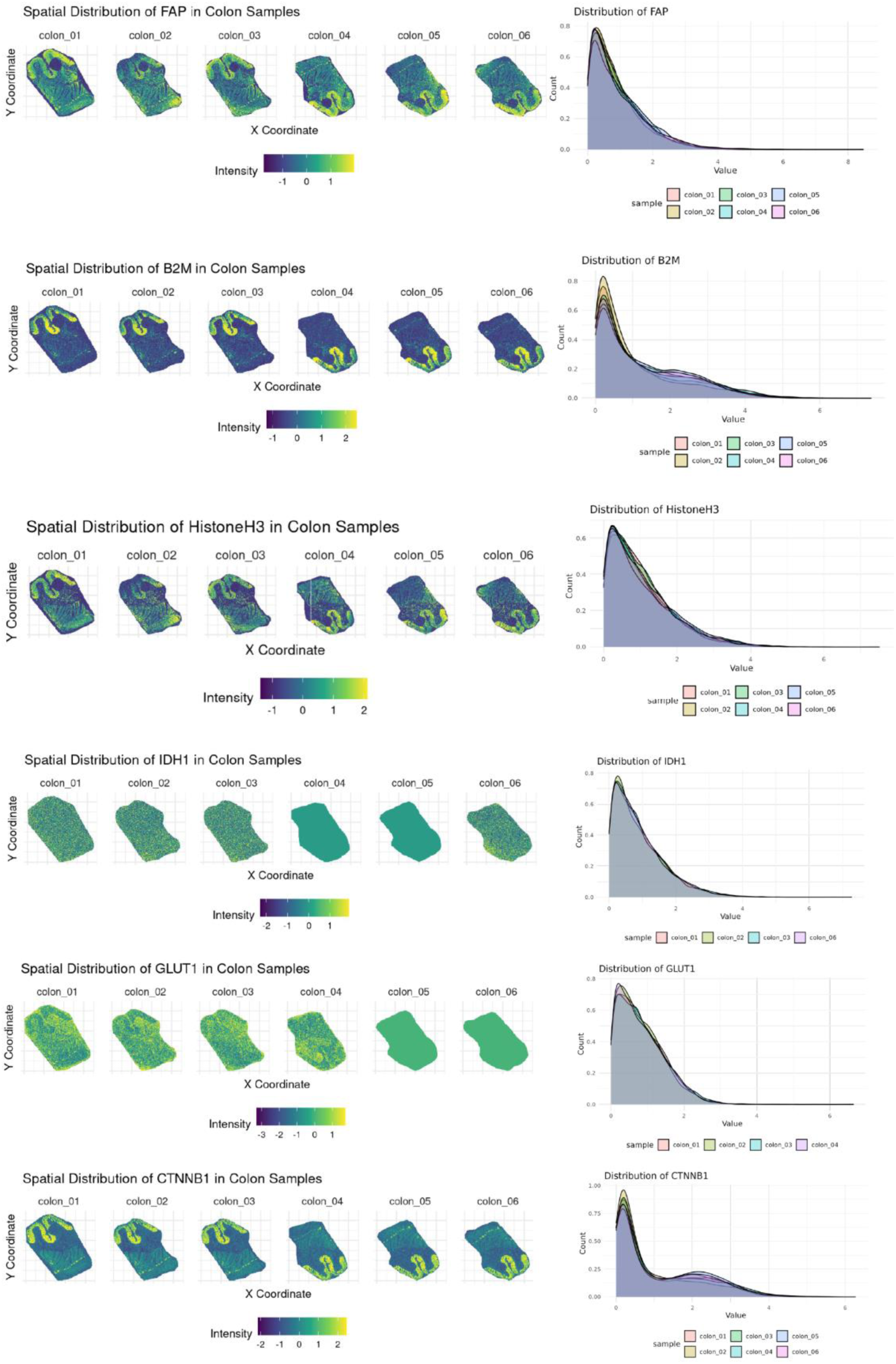
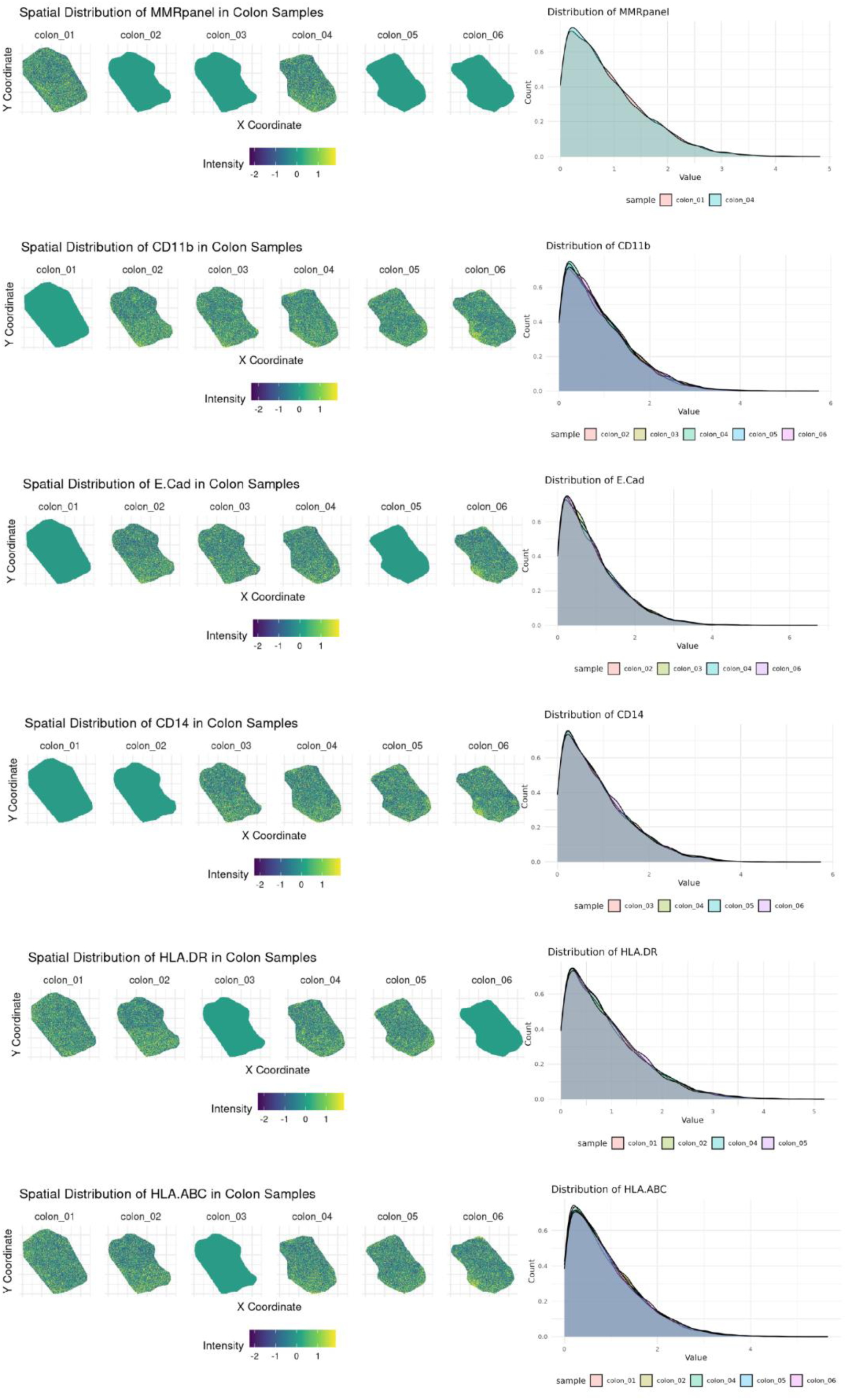
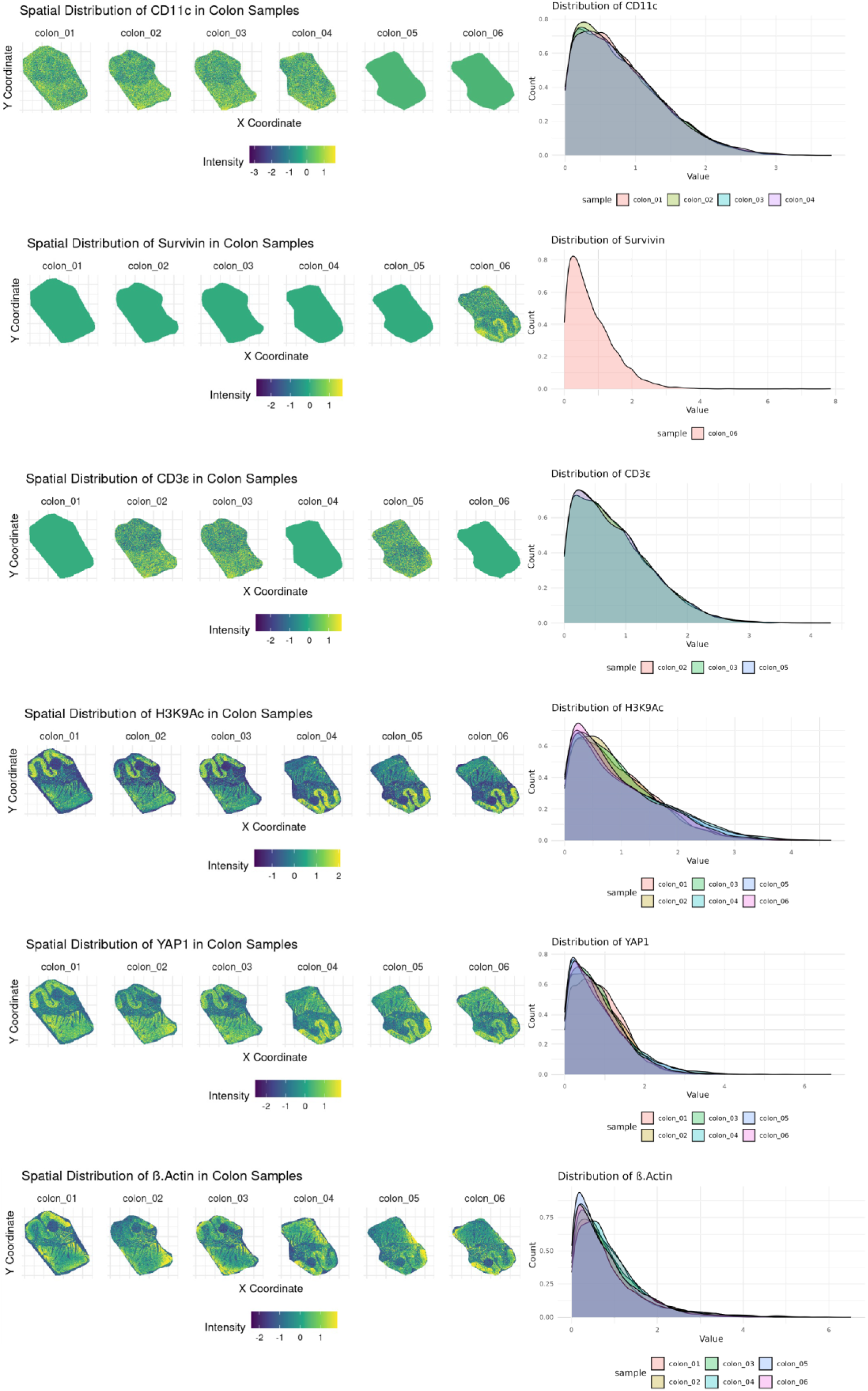
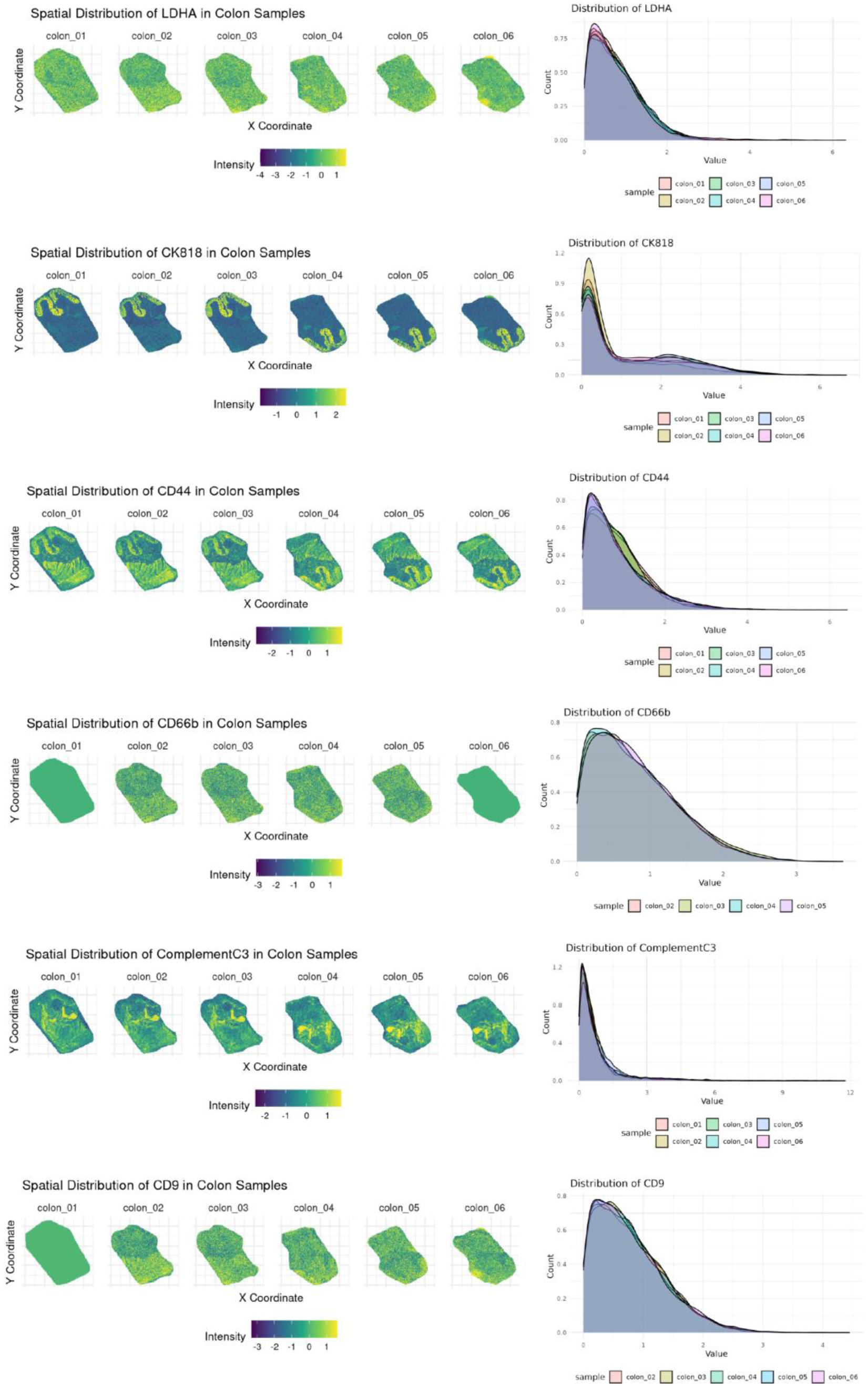
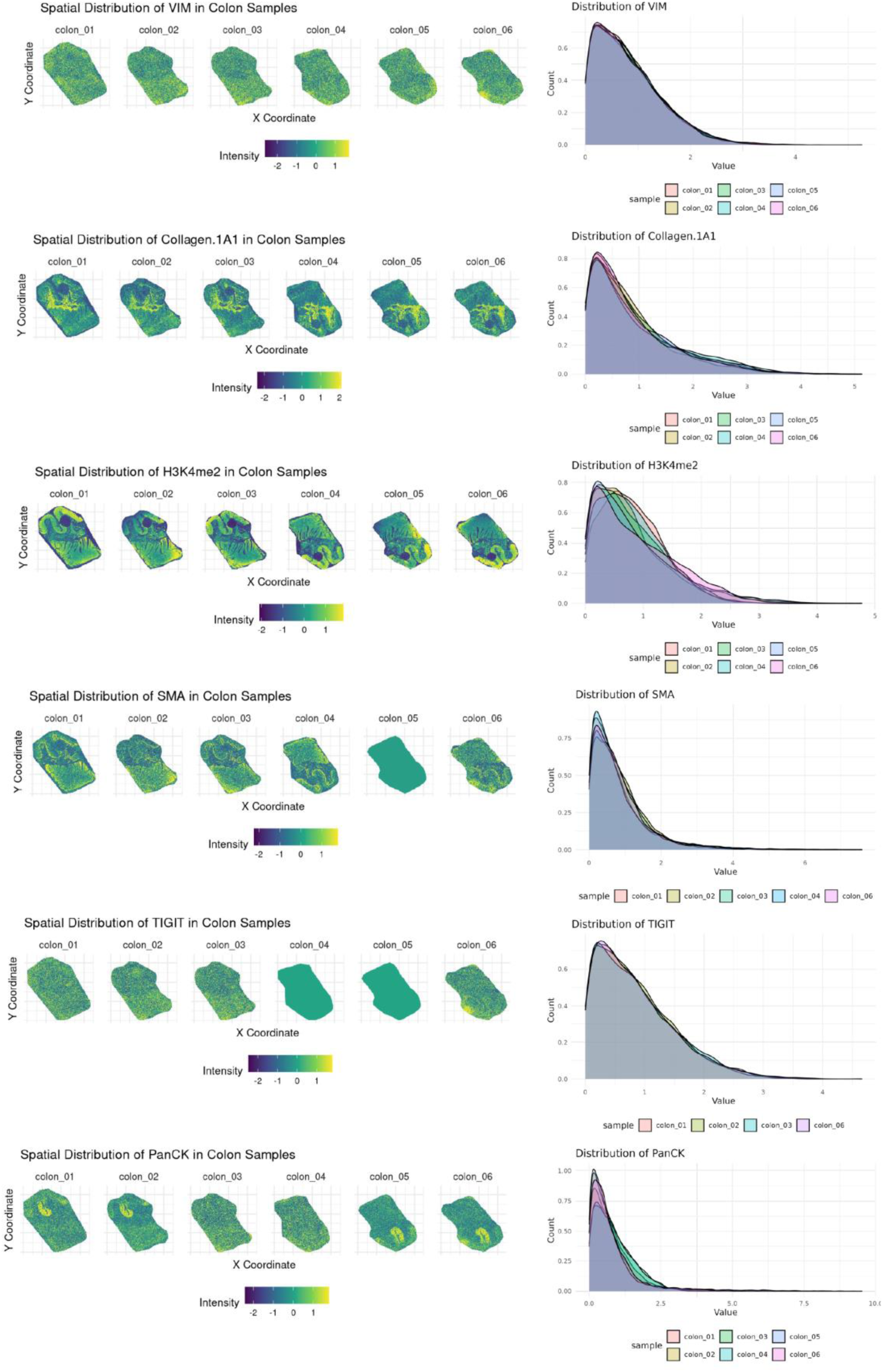
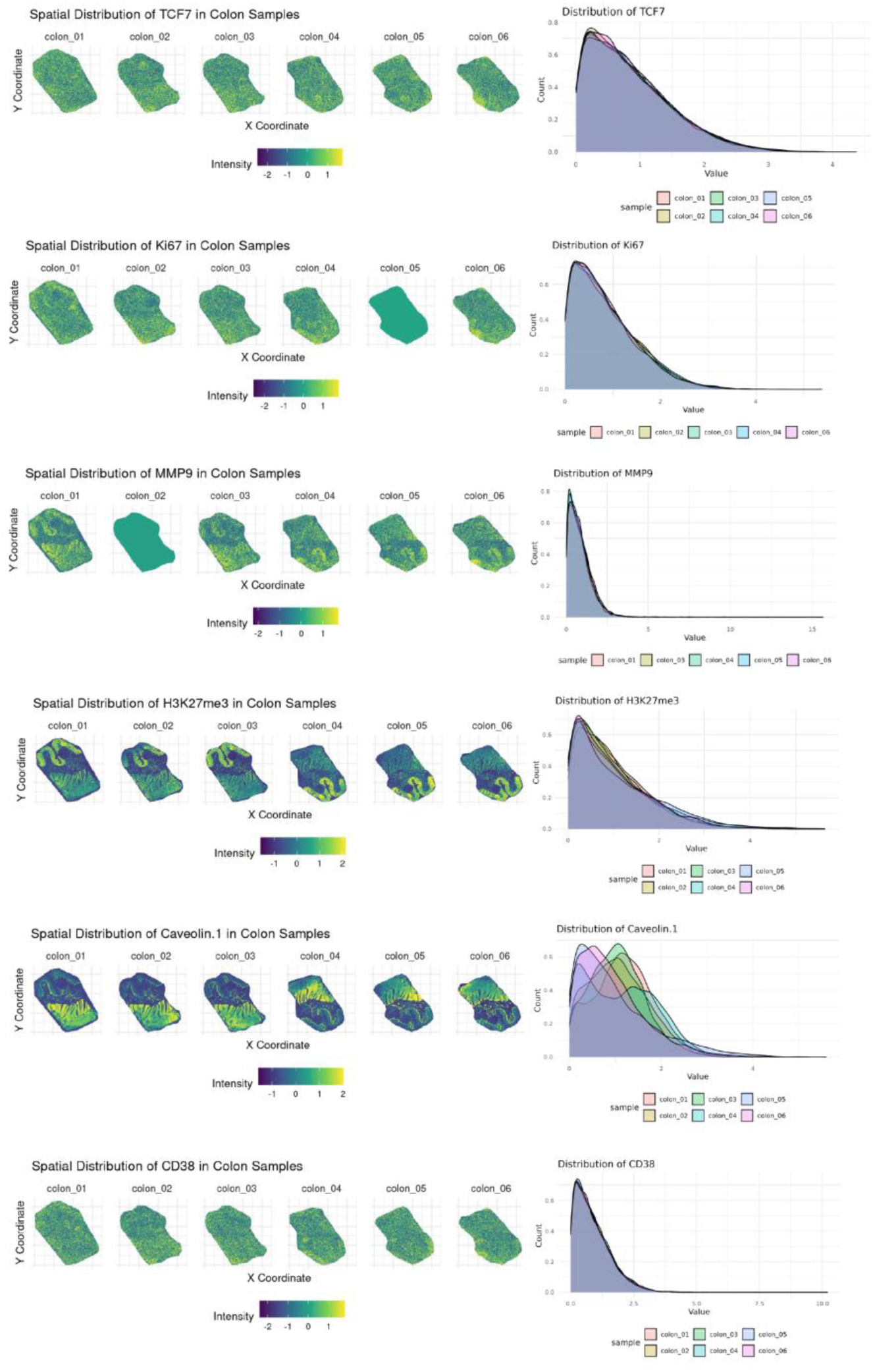
Reproducibility assessment of MALDI-IHC data. (**a, b**) Violin plots show mean expression of all markers across replicates of tonsil (**a**) and placenta (**b**). There is no significant difference between replicates. Horizontal lines link data points representing the same markers. (**c**) The distributions for all markers in the six colon replicates are shown. The ion images (left) show marker intensity, the intensity/density plots (right) show distributions across the entire imaging area, with each colon replicate sample indicated by color.

**Supplementary Figure 5:**
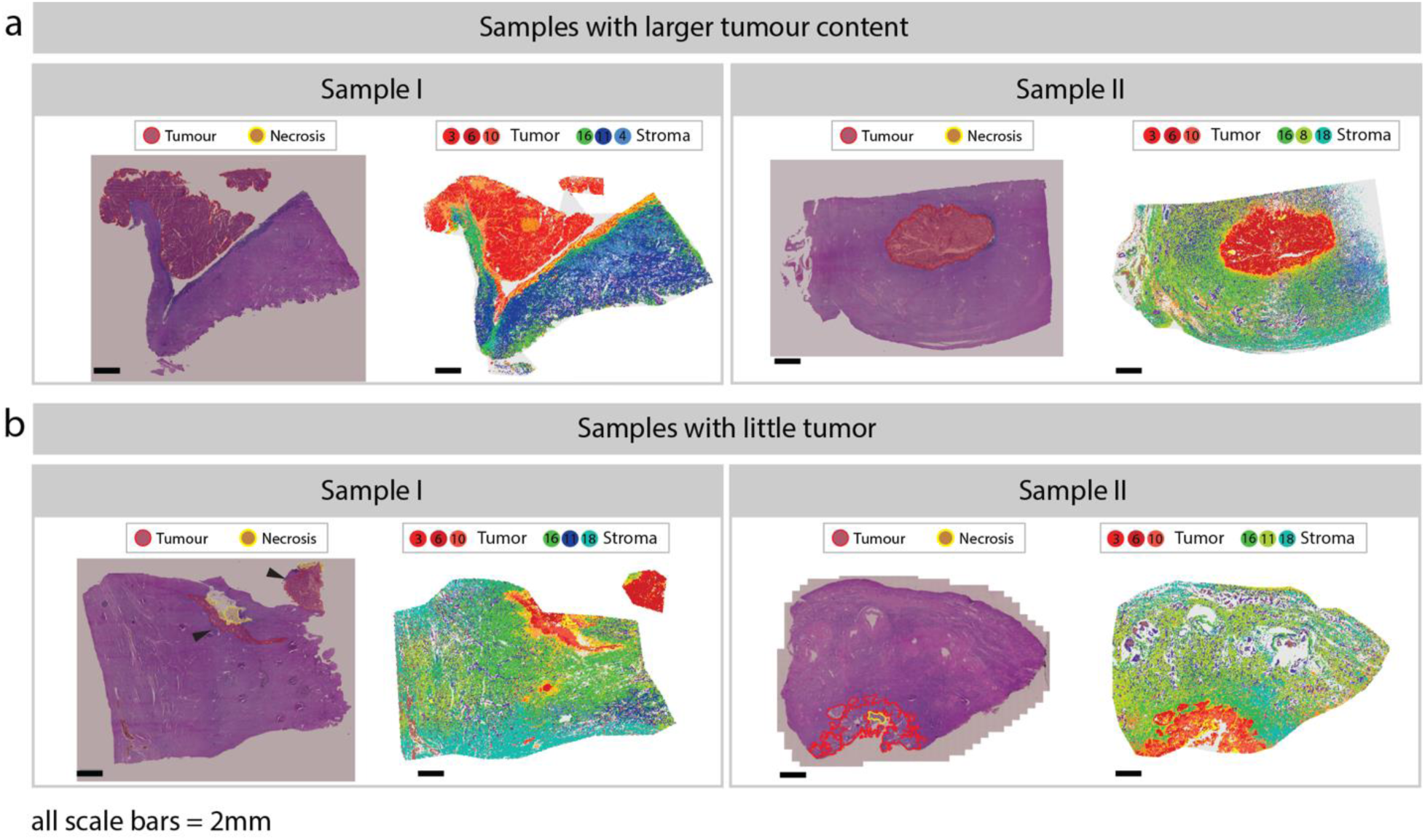
Tumor content varies across samples in the endometrial cancer cohort. **(a, b)** Pathologist-annotated H&E images and the corresponding MALDI cluster map are shown pairwise for exemplary samples with larger (**a**) or smaller (**b**) tumor content. Mapped clusters are indicated by colour. Pathologist annotations were conducted independently, outlining the tumor (red) and necrotic (yellow) regions.

**Supplementary Figure 6:**
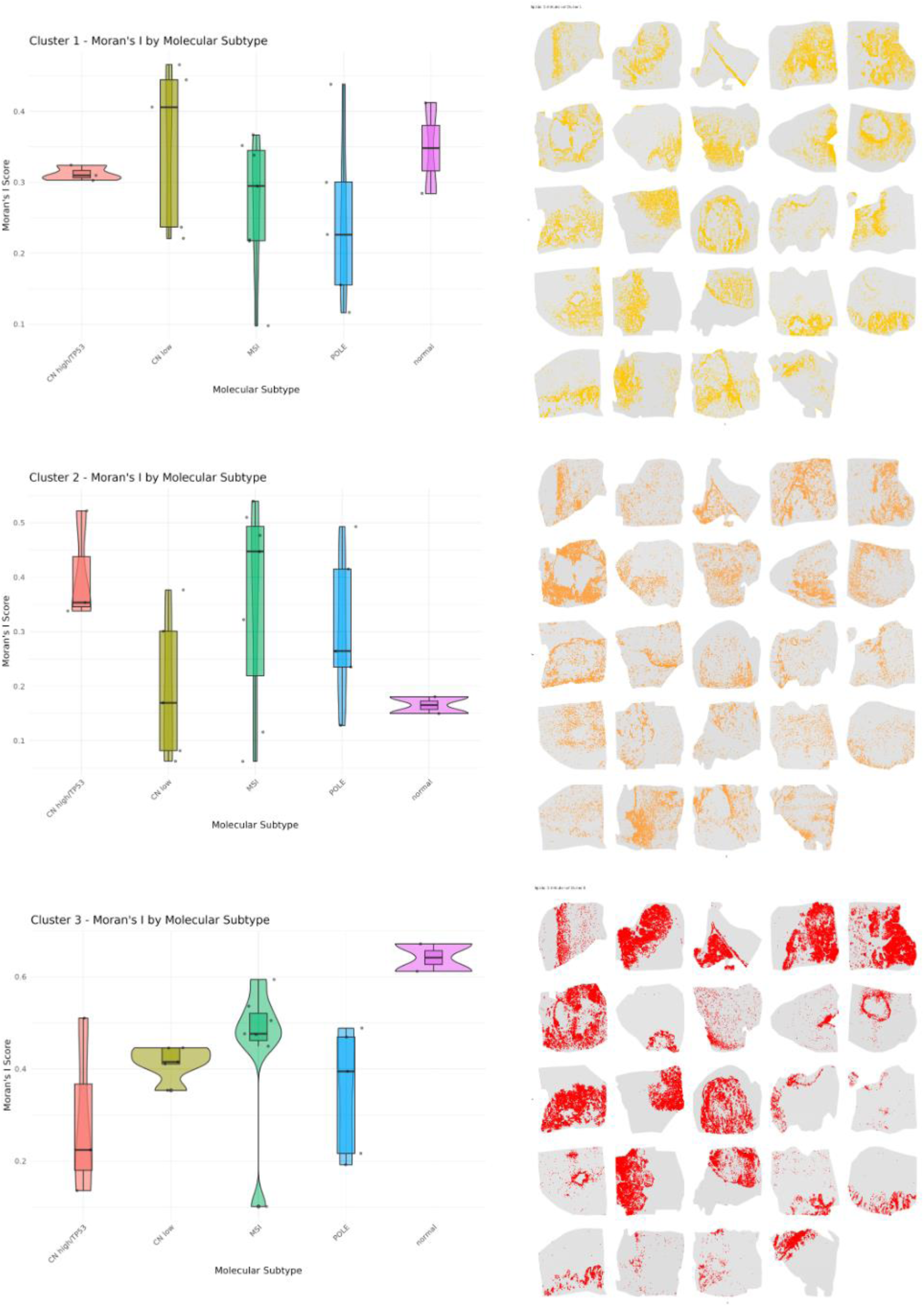
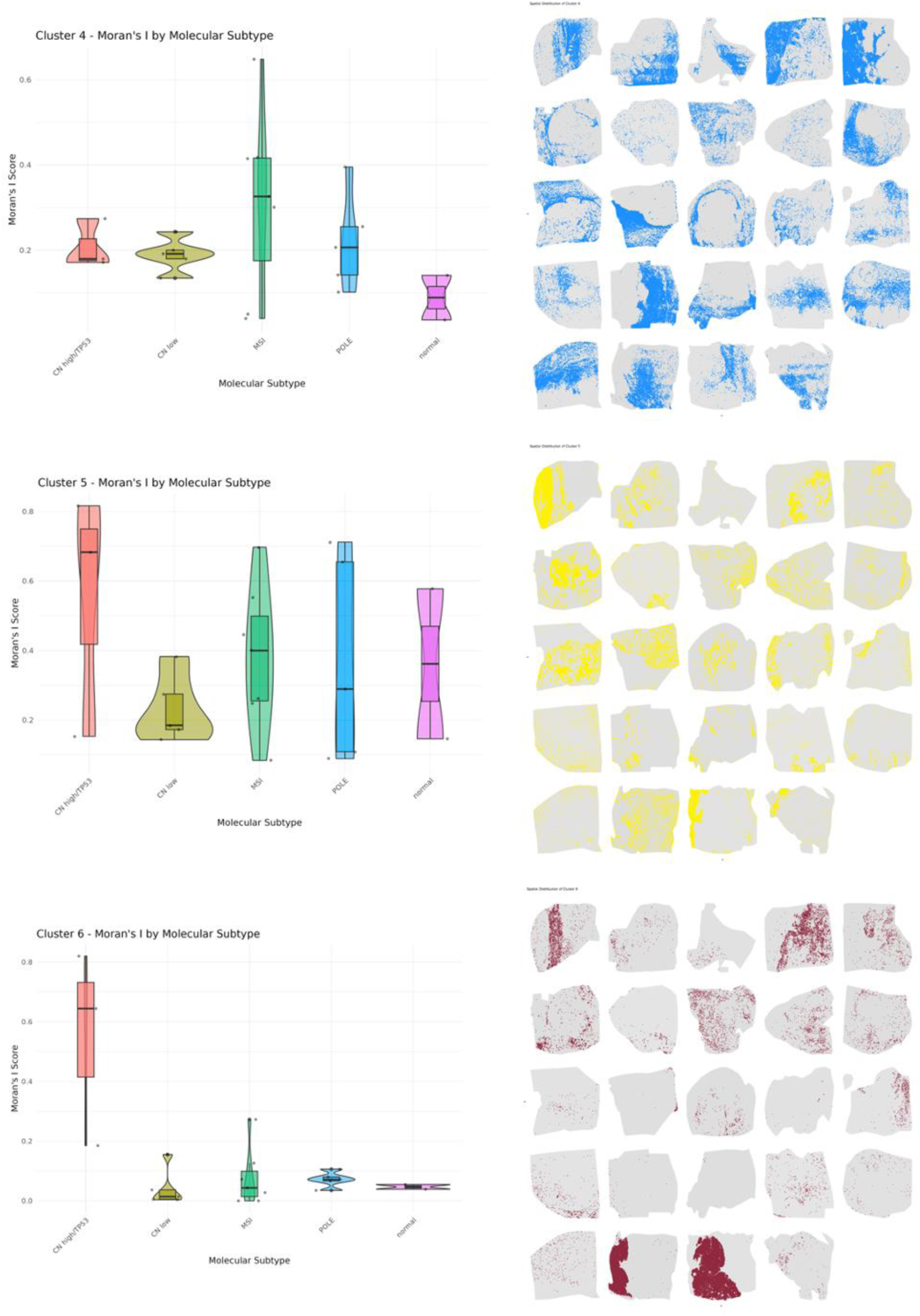
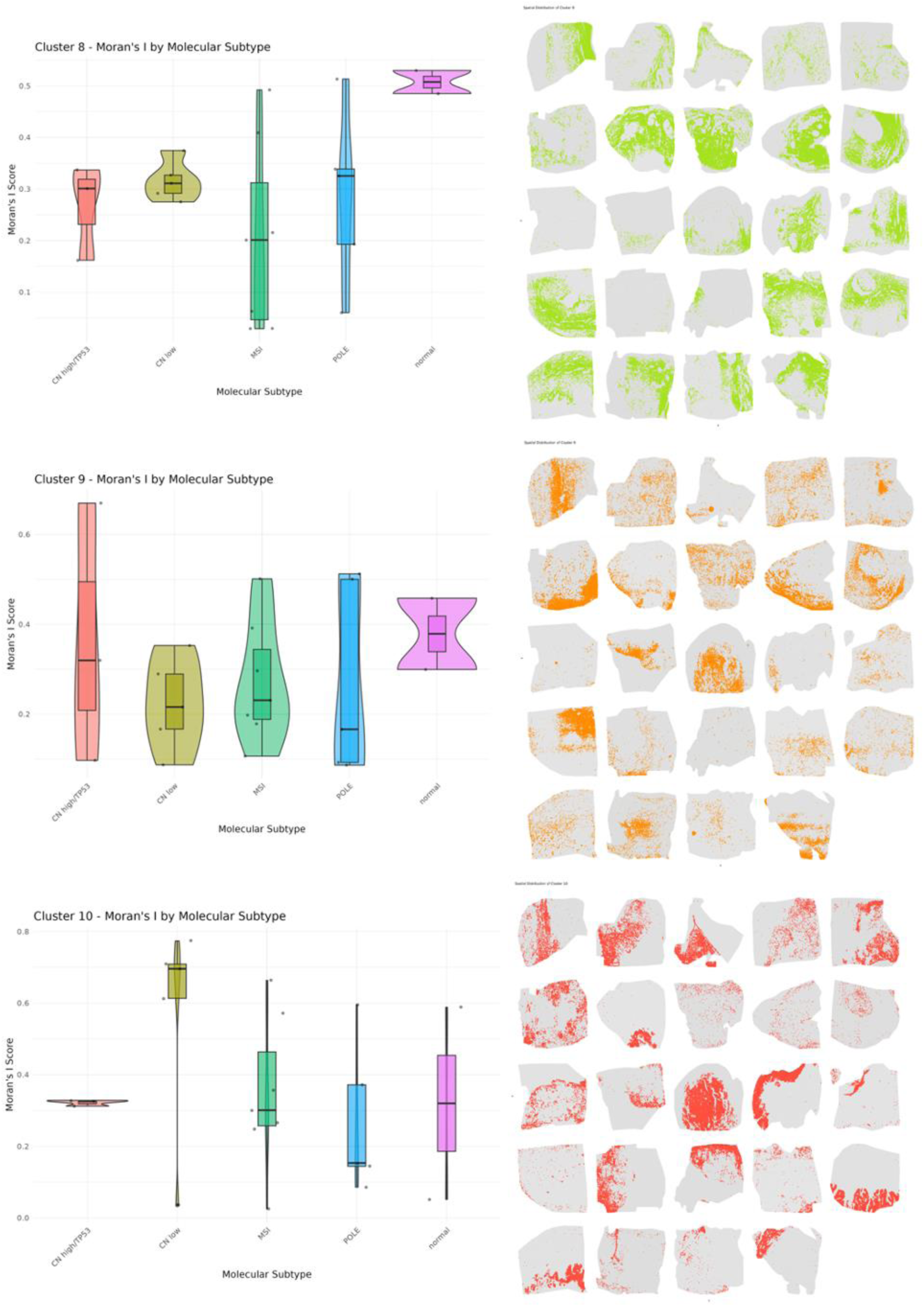
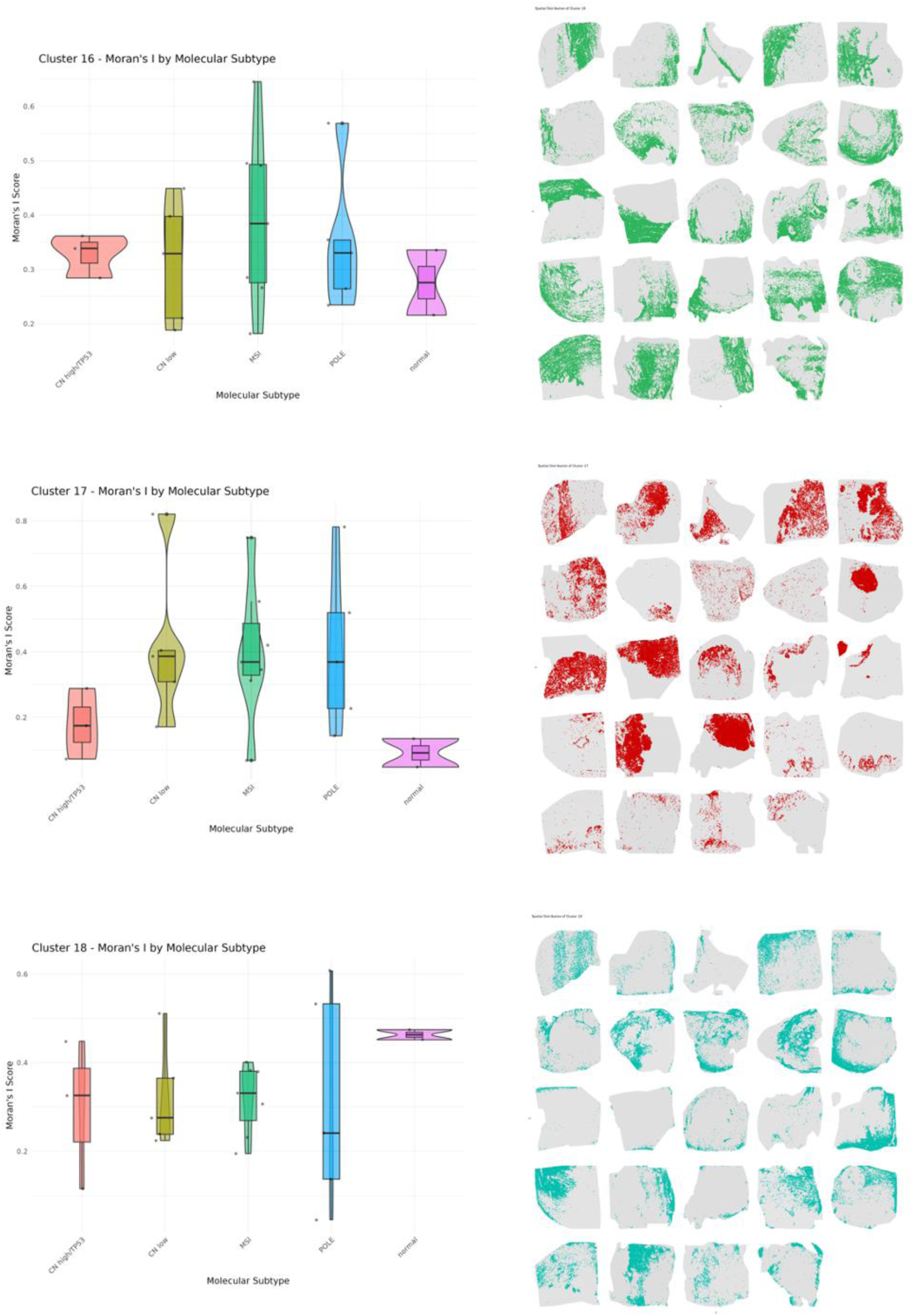
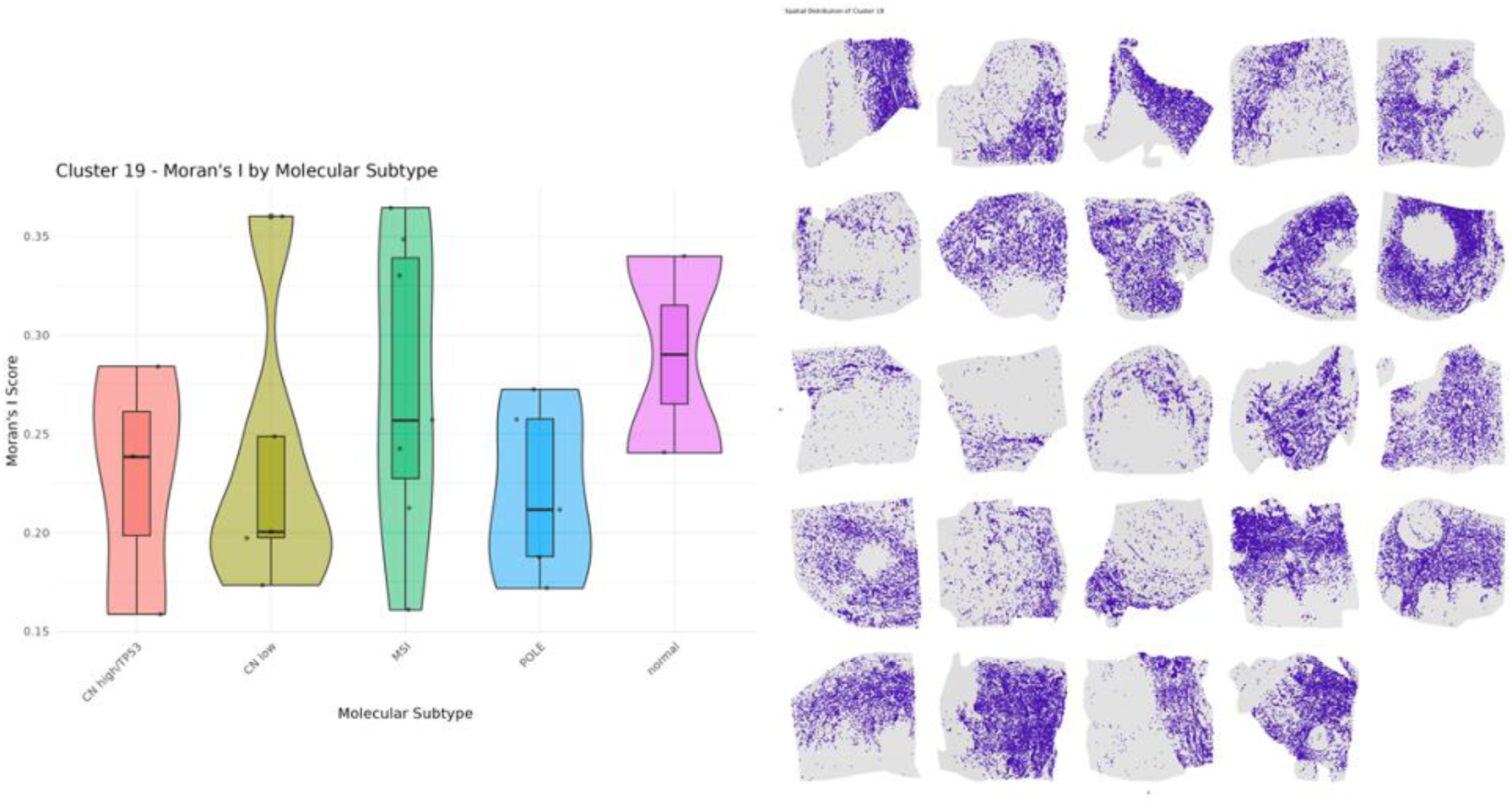
Spatial analysis of segmented regions in the endometrial cancer cohort. For each cluster, shown are violin plots depicting the distri}bution of Moran’s I statistic across molecular subtypes (left) and spatial patterns in all high-quality samples (right).

